# ProteoBridge Connects Proteomics Quantification Outputs to Reusable Analysis Workflows

**DOI:** 10.64898/2026.11.23.768141

**Authors:** Witold E. Wolski, Robbe Devreese, Robbin Bouwmeester, Henry Webel, Leonardo Schwarz, Caroline Jachmann, Sam van Puyenbroeck, Christian Panse, Hubert Rehrauer, Lennart Martens, Marie Locard-Paulet

## Abstract

Quantification tools for bottom-up label-free proteomics, including DIA-NN, Spectronaut, MaxQuant, and FragPipe, use different table layouts, column headers, modification notations, and parameter files. Downstream applications maintain separate readers for these outputs. ProteoBridge lets an application request the same quantity, such as unnormalized ion intensity, across reports from different quantification tools. JSON parsing rules describe each tool’s tables, and software-specific parameter parsers read the settings. The rules currently cover 14 tools and are tested on public inputs. A shared parser organizes the reported quantities into one level for each feature type the tool reports (fragments, ions, peptidoforms, peptides, or proteins) and writes the result as AnnData/MuData, Parquet, or DuckDB. A JSON summary records the structure, provenance, and conversion warnings, such as unmapped modifications. Applications can use the apb-catalog API in Python, R, or TypeScript to map the requested quantity to the field that holds it for each tool and version. Researchers can supply a sample table to add the experimental design and a FASTA database to check peptide sequences. ProteoBridge can serve as a common interface for reading quantification results across tools and versions.

## Introduction

Quantification software for bottom-up label-free proteomics by mass spectrometry (MS), such as `MaxQuant`, `DIA-NN`, `Spectronaut`, and `FragPipe`^1–6^, or the open-source `AlphaDIA`, `Sage`, and `Proline`^7–9^, each has its own table layout and its own notation for modifications and settings. Each downstream application therefore needs a reader that identifies the features and selects the abundances and q-values used for quantification and filtering. For example, the benchmarking platform `ProteoBench`^10^ has its own readers, as have `prolfquapp`, `ProteoPy`, `msmu`, and `AlphaPeptTools`, which analyze quantification results, for instance for differential expression analysis^11–15^. The same table may therefore be parsed differently by each of them. Their readers need updates whenever a tool changes its output, as `DIA-NN` did between versions 1.8 and 2.0, and their developers update them independently. In the `prolfqua` ecosystem, readers of quantification tool outputs account for 16% of the code and 17% of the commits from 2018 to October 2026. In `ProteoBench`, the parsers for parameter files and result tables account for 32% of the code and 26% of the commits from 2023 to October 2026.

This duplicated work is what the Proteomics Standards Initiative (PSI) and many converters set out to avoid. PSI defined standards: mzML for raw data, mzIdentML for identification results, mzQuantML and the tab-separated mzTab for quantification^16–19^. Its newest standard, mzPeak, stores raw data in the Parquet file format and keeps columns outside its schema under an `opt` prefix, bound to PSI-MS terms where they exist^20–22^. Few quantification tools write mzTab, among them `quantms`^23^. Converters handle one kind of data: `msconvert` converts raw files from manufacturer file formats to mzML^24,25^.

Instead of converting files, `mzAPI` proposed for raw MS data to standardize access, with one programming interface over the reading libraries of the instrument manufacturers. Its authors argued that standards written in XML, such as mzML, suit large numeric data poorly, and that export to a common standard keeps only the “lowest common denominator” of each instrument’s data^26^. The manufacturers distribute their own reading libraries, such as Thermo’s `RawFileReader`^27^ and Bruker’s library for timsTOF files. Most quantification tools, among them `DIA-NN` and `AlphaDIA`, build these libraries in directly, while `ProteoWizard`^24,25^ provides one interface over open standards and manufacturer file formats in C++, and the `Spectra` package with its `MsBackend`^28^ one in R (Section S3). Both expose a common set of spectrum variables, a subset of what the raw file records. Native files keep every value, but `mzAPI` left open how a common interface would expose what only one instrument records.

Result tables need no manufacturer library, because quantification tools write them mostly as text tables or Parquet files, but they have no common standard or interface either: which column holds which abundance or q-value (per run or global) differs between tools and versions. ProteoBridge, the AnnData Proteomics Bridge (APB), combines the two routes. Like the converters, it converts the tables of every quantification tool into one annotated data matrix per level, with ProForma identifiers. The annotated data matrix, a matrix of quantities with tables that annotate its features and samples, has represented quantitative omics data for more than two decades: Bioconductor’s ExpressionSet and its successor SummarizedExperiment implement it in R^29,30^, AnnData in Python^31^, and Multi-AssayExperiment and MuData group several such matrices^32,33^. ProteoBridge thus adds no new data model, standard, or file format. Unlike the converters, it keeps every information-carrying column under a name taken from the tool’s header (a column with a single value is kept, with that value, in the provenance record), and it answers the question `mzAPI` left open with a catalog that maps a term, such as unnormalized ion intensity, to the column of each tool and version when a result is read. This design aligns with the division of expertise that Bakunin described with his bootmaker (Section S1). Applications query the catalog through a configurable application programming interface (API) in Python, with ports to R and TypeScript. Climate science reads data this way: `cf xarray` finds a variable by its Climate and Forecast (CF) standard name^34,35^.

The architecture and ideas driving the development of ProteoBridge are summarized in Figure 1, which is organized by three actors, the people who work with ProteoBridge (see Section S2 for the description of all terms used in the figure). The quantification software developer, on the left, knows what the tool writes for its users, from the developers themselves, who improve feature detection, to mass spectrometrists, who might use the outputs to fine-tune their chromatography. Therefore, the tools write many columns, and their export options decide which. We assume that each column is written for a good reason, so ProteoBridge keeps every column it parses. At the peptidoform-ion level, ProteoBridge currently parses up to 70 columns of a `DIA-NN` report^36^. The researcher can provide the experimental design, which `apb2 annotate`, from ProteoBridge’s converter package `apb2`, adds to the ProteoBridge result as the sample table.

**Figure 1:**
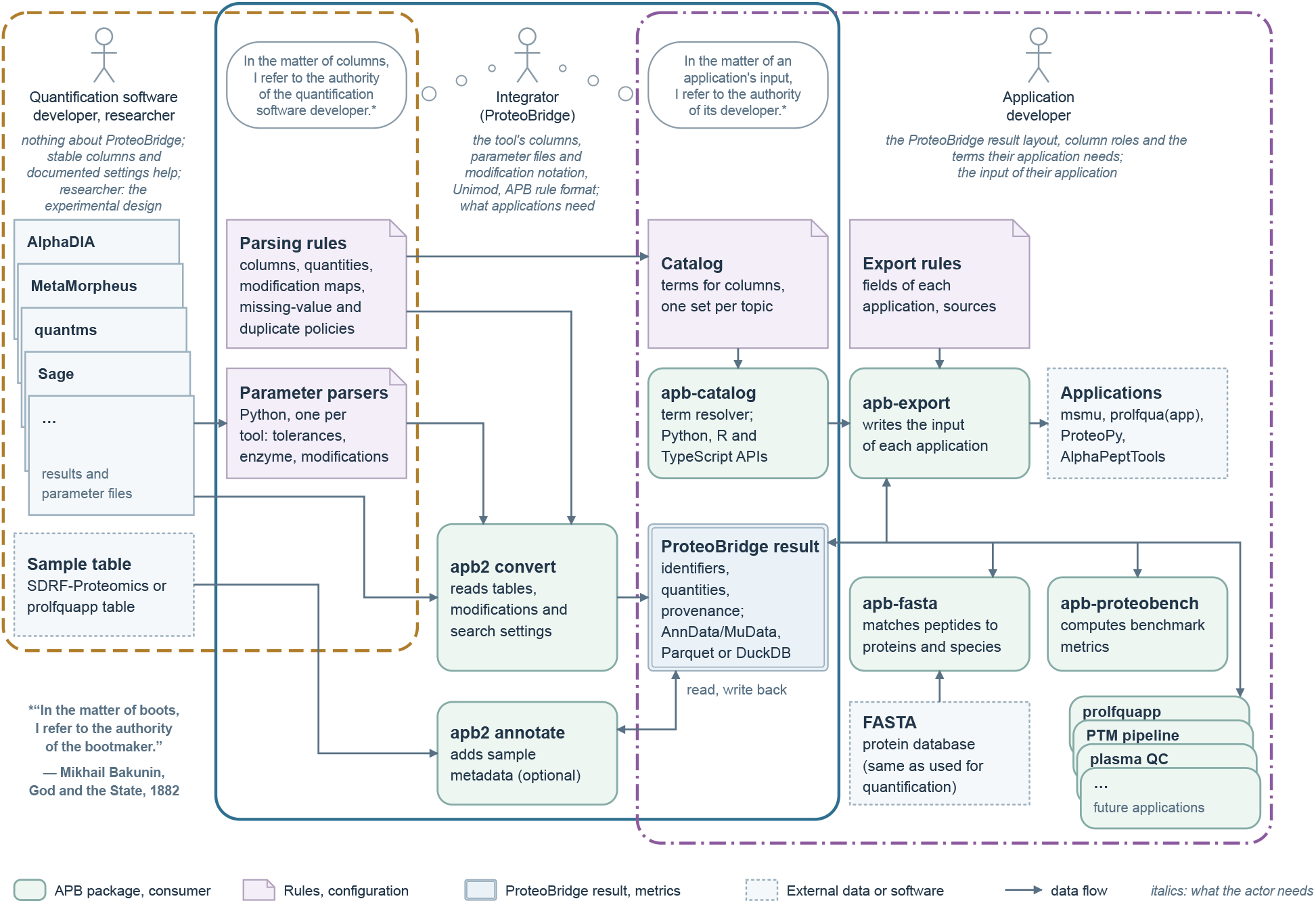
ProteoBridge packages and actors. Quantification software writes result tables and parameter files, and researchers supply the sample table. Integrators write and test the parsing rules, with their modification maps, and the parameter parsers. From these, `apb2 convert` builds the ProteoBridge result: feature identifiers, quantities, and provenance. `apb2 annotate` adds sample metadata from the sample table. `apb-catalog` keeps its terms in sets, one per topic, for example abundance or confidence, and maps each term, for instance unnormalized ion intensity, to its corresponding column for each quantification tool and version. Consumers read and optionally enrich the result: `apb-fasta` adds protein and species matches when a FASTA file is provided, `apb-proteobench` adds benchmark metrics when applicable, and `apb-export` applies the export rules to write input for applications. The two speech bubbles adapt the bootmaker quotation below the first region; Section S1 explains it.

The application developer, on the right, knows the application and defines a minimal interface for it: the data that the application needs to be fully functional. This follows the interface segregation principle of software design: a client should depend only on the parts of an interface that it uses^37^. The fewer terms an application requires, the more quantification tools it can read. The developer checks the catalog, `apb-catalog`, for the terms that name these data, for instance unnormalized ion intensity or the aligned retention time of an ion. If a term is in the catalog, ProteoBridge returns its values for every quantification tool that writes them; if it is not, an integrator can add it.

The integrator, in the middle, knows neither every use for which a tool writes its columns nor every idea of the application developers, but brings the two sides together. To do so, the integrator needs to know the documentation and column names of the quantification tools and the annotated data matrices of the ProteoBridge result. With this knowledge, the integrator writes and tests the parsing rules and parameter parsers, from which `apb2 convert` builds the ProteoBridge result, and fills the catalog, linking each term of the application developers to the field of the ProteoBridge result that holds it for each tool. Other actors can take on part of this work: quantification software developers can contribute the parsing rules and parameter parsers for their tools, and application developers the catalog entries for the terms they need.

Existing projects read the tables of quantification tools into one data model or standard (Section S3): QPX defines a standard, stored in Parquet, for exchanging quantitative proteomics results, while `AlphaPeptTools`, `msmu`, and `ProteoPy` read results into AnnData or MuData^13–15,38^. Each maps a tool’s columns to fields of its own, mostly with one quantity per level, and drops the other columns or reads them only on request. `psm utils`, the closest to ProteoBridge in design, does the same for the peptide-spectrum matches of many search engines^39^, and `SWATH2stats` converts SWATH-MS results for several statistical applications^40^. ProteoBridge, in contrast, keeps the columns its parsing rules read under the tool’s own names and lets applications select them through the catalog; with this strategy, it maps many quantification tools to many applications: it currently reads the output of 14 quantification tools^36^ and writes ten export targets.

## Experimental section

### Input data

We selected the public datasets for the conversion measurements in November 2026. The 198 `ProteoBench` submissions, each containing a result table and parameter file, span seven quantification modules (Table S3)^10^. Five large exports come from the benchmark inputs of `directLFQ`^41^ and `MSstats+`^42^. The `MaxQuant` dataset holds 200 HeLa replicate runs^43^, and the `Spectronaut` dataset covers the four-species LargeFC benchmark^44,45^. The `MSstats+` reports cover a K562 dilution series (`DIA-NN`, 40 runs), a mixture of human, yeast, and *E. coli* proteomes (`Spectronaut`, 18 runs)^46^, and the Pan-Cancer Proteome Atlas (`DIA-NN`, 1,236 runs)^47^. Additional examples use public test data from the same sources. The export example uses `ProteoBench` submission 0132a468 to the benchmarking module for data-independent acquisition (DIA) with all-ion fragmentation (AIF), a `DIA-NN` 1.9.2 report of six runs, with the module’s FASTA and sample table in SDRF-Proteomics format^48^ from `apb-proteobench`. We downloaded these inputs and ran the workflows with our test harness, `APB Studio`, which uses `Snakemake`^49^. Section S4 summarizes the datasets and provides their raw-data accessions.

### Conversion cost

We converted each input twice and wrote AnnData/MuData (HDF5): once at the peptidoform-ion level only and once at every level that the input reports, one conversion at a time with four `Polars` threads. Conversion time includes compilation, reading, and parsing, but not writing the result, and peak memory is the process’s maximum resident set size. Section S5.1 describes the hardware and the measurement details.

### Storage formats

We converted 19 inputs to AnnData/MuData (HDF5), Parquet, and DuckDB: 12 `ProteoBench` submissions, a `MaxQuant` export (PXD070049), a `MetaMorpheus` export (PXD003881), and the five large inputs (Section S4). Output size is the on-disk size of the stored result. For Parquet, it is the summed size of all files in the Parquet directory. Output size excludes the JSON and timing files written next to each output.

### Implementation

ProteoBridge and its companion packages are written in Python, building on Polars, NumPy, SciPy, pandas, pydantic, cyclopts, and loguru^50–56^.

### AI-assisted development and verification

We wrote much of ProteoBridge’s code and parsing rules with AI coding assistants. An agent skill is an instruction file that an assistant loads for a task. Because most tasks we gave the assistants were complex, we first wrote detailed planning documents, which the assistants drafted in a dialogue with us through the skill `interview-to-spec`. To keep the code modular, we set design rules with the skill `directed-folder-imports`, and to keep it readable and reviewable, with the skill `polymorphism-over-discrimination`.

We validate ProteoBridge with continuous integration and a test harness. On every change, GitHub continuous integration runs `Ruff`^57^, strict `Pyright` type checking^58^, unit tests with branch coverage, and a documentation and package build per ProteoBridge package. `APB Studio` runs workflows such as conversion, `ProteoBench` scoring, and export on public inputs (Input data). From each dataset’s run log, `APB Studio` flags failed steps and plots the runtime and peak memory of all steps, including reading, converting, and writing, against input size. Because most `ProteoBench` modules mix human, yeast, and *E. coli* proteins in known ratios, the metrics it displays show whether a conversion kept the quantities intact. From the JSON summary of a ProteoBridge result, `APB Studio` shows the columns and value distributions of the `var`, `obs`, and `varm` tables. It also lists the warnings that each step recorded (Section S6). The conversion step records modification notations missing from the modification map, unreadable numeric cells, feature columns whose values differ between rows of one feature, and the empty or single-valued columns and layers it left out.

## Results and Discussion

While converting, ProteoBridge sorts the quantification tool’s data by level: each value goes to the level of the feature it describes, which can be a peptidoform fragment (a fragment ion of a peptidoform ion), a peptidoform ion (a peptidoform with its charge), a peptidoform (a peptide sequence with its modifications), a peptide, or a protein. Each level is a matrix of quantities, with a feature table that annotates its rows and a sample table that annotates its columns. Within a level, a value that varies between samples goes into a matrix, and a value that describes only the feature goes into the feature table. Sorting the data this way documents and validates the tool’s output: the structure states what each value describes and what it depends on, and a value placed in the feature table must be the same on every row of its feature, or ProteoBridge records a warning.

Each level of the ProteoBridge result is an annotated data matrix with the parts of AnnData^31^: a sample table (`obs`), a feature table (`var`), feature-aligned tables, for example FASTA matches (`varm`), the primary abundance (`X`), and further quantity layers. The shared parser builds the identifier columns of three levels: `ProForma ion` for peptidoform ions, `ProForma peptidoform` for peptidoforms, and `ProForma peptide` for peptides. These columns follow ProForma 2.0 notation with Unimod modifications ^59,60^, and the PSI-MS controlled vocabulary has a term for each, such as MS:1003270 for peptidoform ions^22^. The parser assigns two roles to feature-table columns: `protein assignment` for the column that holds the protein group, and `fasta accessions` for the column of FASTA accessions. Abundance layers get the `abundance` role, so consumers can find columns and layers by role instead of by software-specific names.

A `DIA-NN` report shows why we chose the annotated data matrix. The report has one row per precursor and run, and repeats the protein-group q-values `PG.Q.Value` and `Global.PG.Q.Value` on every row of the group’s precursors. In the ProteoBridge result both sit at the protein level, the per-run q-value as a layer and the q-value across all runs in the feature table.

Besides every result, ProteoBridge writes a JSON summary. For each level, the JSON lists the sample and feature tables with their columns and types, the layers with their dimensions and missing values, and the roles. It holds one provenance record per step that wrote the result. The record of the conversion, for instance, keeps the parsing rule and each input file with its checksum. Each summary entry counts one kind of problem, such as unreadable cells; a count above zero needs attention. We chose to record any problem that leaves the result usable, such as an unmapped modification, as a warning. The conversion stops only where the result would be wrong, as when a required column is missing.

apb2 convert reads the output table and, optionally, the parameter file, using the parsing rule corresponding to the quantification tool and its version, and converts every level the tool reports or one selected level.

A parsing rule is a JSON file that describes one output layout of a quantification tool, for a range of its versions. `DIA-NN`, for example, currently has three rules, selected by the software version recorded in the parameter file. Each rule selects columns that identify samples (run or file name) and features, such as the modified peptide sequence and charge of a peptidoform ion. It assigns each other column to a matrix or to the feature table. A rule names each layer and column after the tool’s header, with two changes: a wide layout writes one column per sample, such as `MaxQuant`’s `LFQ intensity S1`, so the layer drops the sample (`LFQ intensity`), and dots and spaces become underscores, so that a name works unquoted in Python, R, and SQL (`Precursor.Quantity` becomes `Precursor Quantity`); the rule records the header that each name comes from. The rule also declares which values count as missing, for example intensities of zero, and how repeated rows for one feature and sample are read, for example a protein quantity that a report repeats on each precursor row. Parsing rules currently cover 14 tools, among them `DIA-NN`, `Spectronaut`^3^, `MaxQuant`, and `FragPipe`. Section S7.1 lists them all, and the apb2 documentation gives their supported versions, file layouts, and levels. Compilation checks the parsing rule against the table header and configures each step from it. The shared parser then runs the same steps in the same order for every tool (Figure 2). Where a tool writes its output in several tables, a preparation step joins them before the shared parser reads them (Section S7.2).

**Figure 2:**
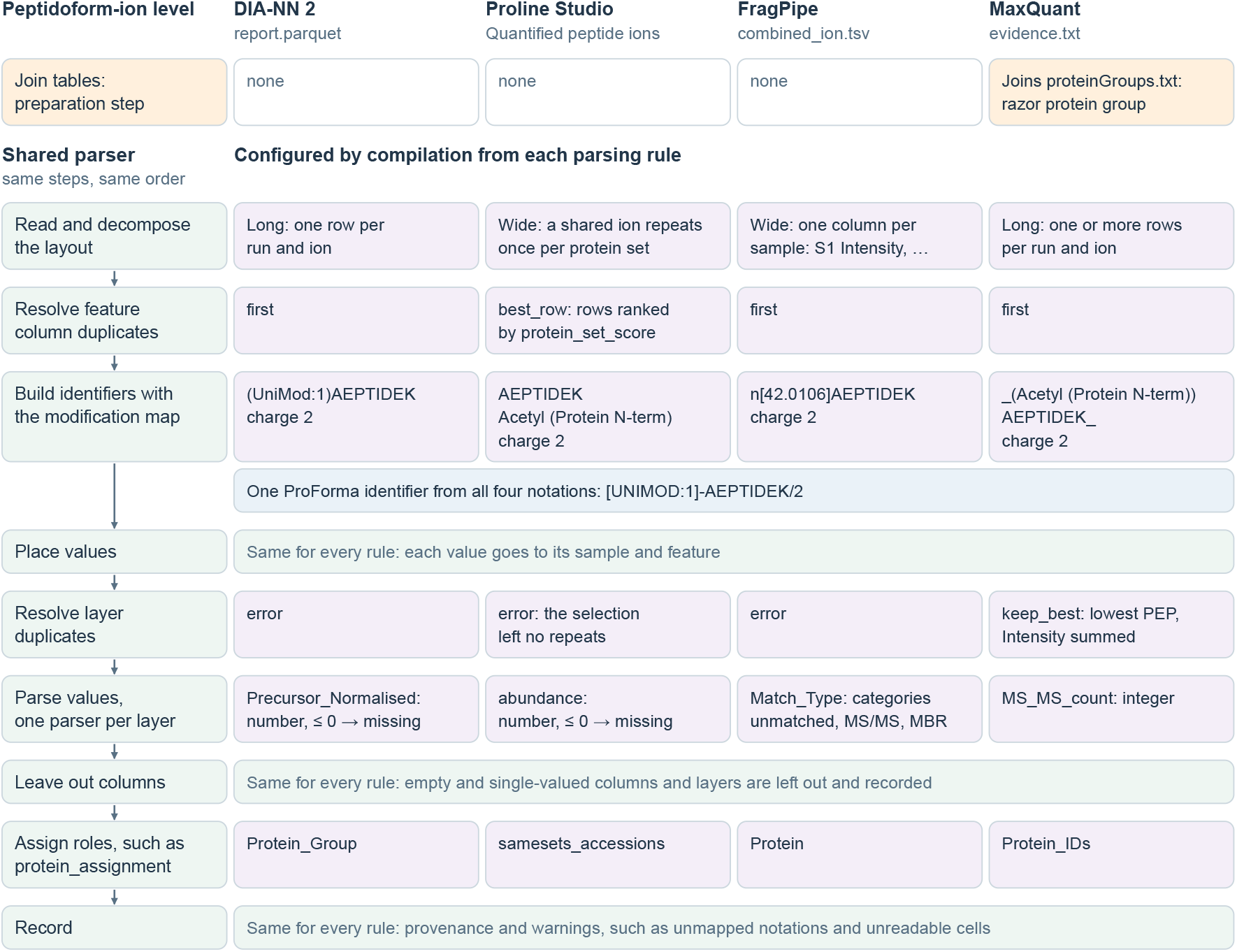
The shared parser at the peptidoform-ion level, for four quantification tools. The steps (green) run in the same order for every tool. Compilation fills in their settings (purple) from the parsing rule of each tool. The orange row is a preparation step that joins the `MaxQuant` evidence and protein-group tables before parsing. Some tools write one feature in several rows: `Proline Studio` repeats a shared ion once per protein set. A feature column, such as the protein accession, then needs one value per feature: `first` takes the first row, and `best row` the best-ranked row. A layer cell, the value of one sample and feature, can repeat as well: `error` stops the conversion, `same` requires equal values, `sum` adds them, and `keep best` takes the best-ranked row. The four tools write protein N-terminal acetylation differently, and all four notations give the same ProForma identifier (blue). Each layer has its own value parser, and the figure shows one per tool. Empty and single-valued columns and layers are left out and recorded.

Where a quantification tool does not write ProForma, its parsing rule declares a modification map, a table that links each tool-specific modification notation to a Unimod accession and its position. The shared parser applies the map. Section S7.3 shows how the tools write three peptidoforms.

The parameter parsers were originally written as part of the `ProteoBench` codebase. Each parsed record holds the software version, acquisition mode, enzyme, missed cleavages, mass tolerances with their unit (ppm or Da), and modifications. Parameter files vary as much as modification notations (Section S7.3). Section S8 reports which settings ProteoBridge recovered from the `ProteoBench` submissions.

We measured conversion on two kinds of public input: 198 `ProteoBench` submissions, which are workflow outputs that users upload to benchmark a quantification tool, and five large exports from the benchmarks of `directLFQ` and `MSstats+`. Figure 3A and B show conversion cost against input size for these 203 inputs. For the 198 `ProteoBench` submissions, median conversion time was 1.33 s at the peptidoform-ion level and 1.50 s for all reported levels, with median peak memory of 0.6 and 0.7 GB. The five large inputs took 6.1 to 154.7 s at the peptidoform-ion level and 7.4 to 627.0 s for all reported levels, with up to 26.0 GB of peak memory. On a separate 150-run subset of the 1,236-run `DIA-NN` report, not part of Figure 3, splitting the packed fragment columns as numbers instead of text lowered the peak memory from 18.0 to 9.3 GB. Besides input size, the numbers of samples, features, and levels to convert affect cost (Section S5).

**Figure 3:**
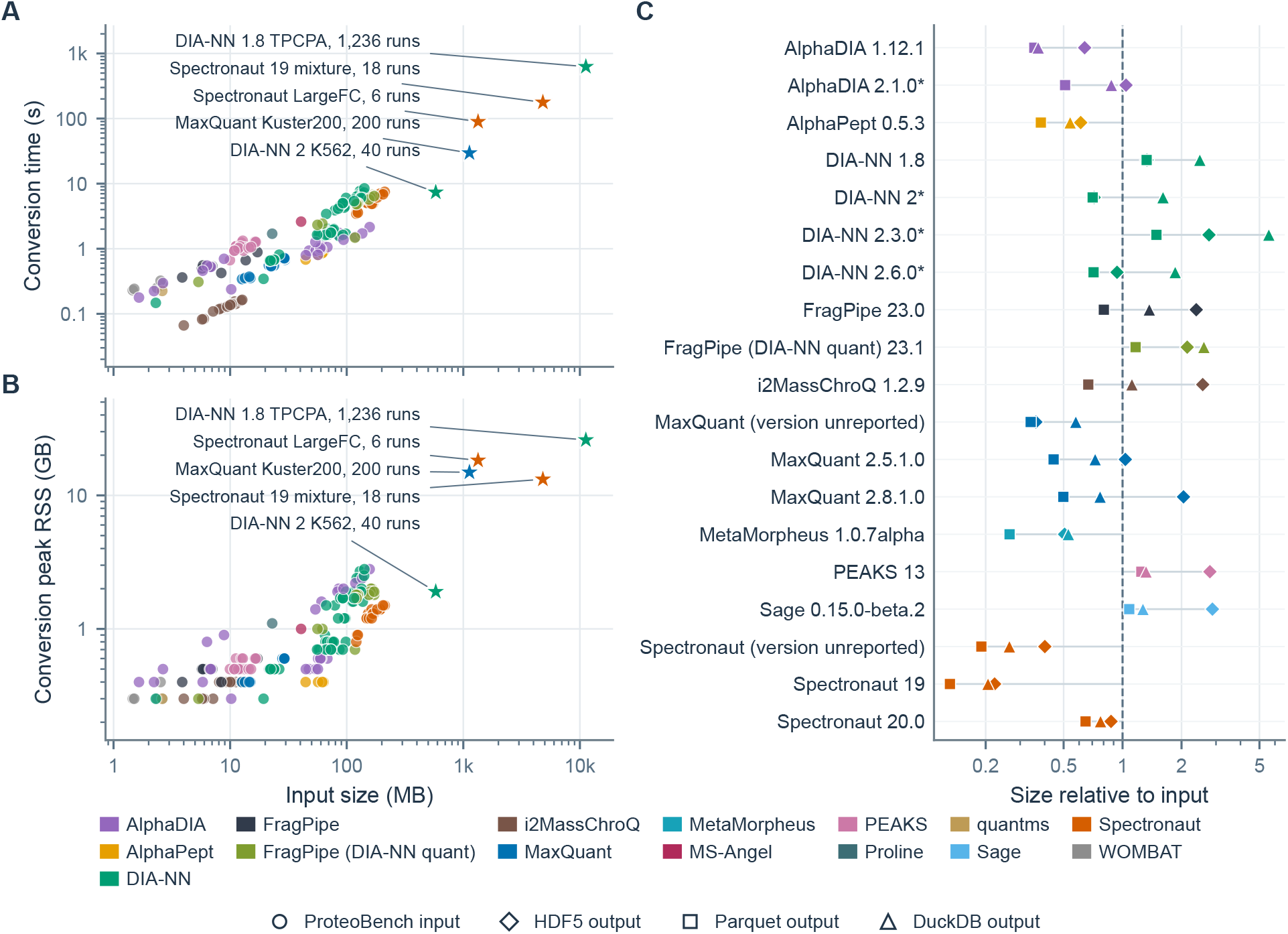
ProteoBridge conversion and storage cost. (A) Conversion time and (B) peak memory (resident set size, RSS) versus input size, for the conversion of 203 inputs to all reported levels, written as MuData (HDF5); the time covers compilation, reading, and parsing, not writing. Circles mark the 198 `ProteoBench` submissions, and stars mark the five large `directLFQ` and `MSstats+` datasets, labeled with dataset and run count. Color identifies the quantification software. Input size is the stored size, gzip-compressed for text inputs. (C) AnnData/MuData (HDF5, diamonds), Parquet (squares), and DuckDB (triangles) output size relative to the size of the same input. The dashed line marks the input size; an asterisk marks an input that is already a Parquet file. Each row lists the tool and the version it reports. All axes are logarithmic.

`apb2` stores the result as AnnData/MuData in HDF5^61,62^, as a Parquet^63^ directory written through `Apache Arrow`^64^, or as a DuckDB^65^ database; Figure 3C compares the three for 19 inputs. Summed over these inputs, the gzip-compressed AnnData/MuData output takes 91.4% of the input size, Parquet 86.0%, and DuckDB 159.3%. For a typical input, Parquet (median 64.7% of the input size) and DuckDB (87.6%) store a result in less space than the input, and AnnData/MuData (103.3%) in about the same. Parquet and DuckDB, however, keep the tables as `apb2` builds them, while AnnData/MuData stores each layer as a numeric matrix; this is why parsing rules map a layer of labels, such as the match type of `FragPipe` (MS/MS, MBR for match-between-runs, or unmatched), to integer codes. A browser can query Parquet directly, and DuckDB answers SQL queries without loading the whole result, which eases the development of web applications. `apb2 reformat result.h5mu result.parquet` converts a stored result between the formats.

`apb2 annotate` adds the experimental design that the researcher supplies, from an SDRF-Proteomics^48^ or `prolfquapp` sample table. It joins the table to the result samples by run name and records the matches.

Quantification tools often report several abundances or q-values for a feature, and an application must know which one it reads. `DIA-NN` reports fragment- and MS1-based intensities, both with and without cross-run normalization, while `MaxQuant` reports intensity, MaxLFQ, and iBAQ for proteins. To distinguish these quantities, `apb-catalog` keeps its terms in sets, one per topic, for example abundance, confidence, and chromatography (Section S9). Under each term, `apb-catalog` lists the column every tool version writes, if any, as a shop catalog lists every supplier’s boots. The catalog thus maps many tool columns to many terms, and an application asks for a term once instead of keeping a reader for each tool. The abundance set describes a quantity by its kind, `intensity` or `ibaq`, and by qualifiers, such as the normalization or the aggregation of lower-level features (Section S9.1). Each term has a key that fixes its kind and qualifiers: `intensity` is an unnormalized intensity, and `intensity norm settings` an intensity with the cross-run normalization set in the search.

Application developers state the terms their applications need, such as unnormalized ion intensity, and mark each as required, recommended, or optional, so an application can keep its interface minimal. An application then asks for a term, not a column (Figure 4): for unnormalized ion intensity, the catalog returns `Precursor Quantity` for `DIA-NN`, `Intensity` for `MaxQuant`, and nothing for `AlphaDIA`: without its run parameters, it is unclear whether its intensities are not normalized, so `AlphaDIA` has an entry only for `intensity norm settings`. A consumer reads each level through the catalog with the interface of AnnData^31^, but names terms where AnnData names columns (Section S9). An application therefore needs no reader of its own for any quantification tool. When the integrator adds a parsing rule and catalog entries for a new tool or version, the application reads its results with unchanged code.

**Figure 4:**
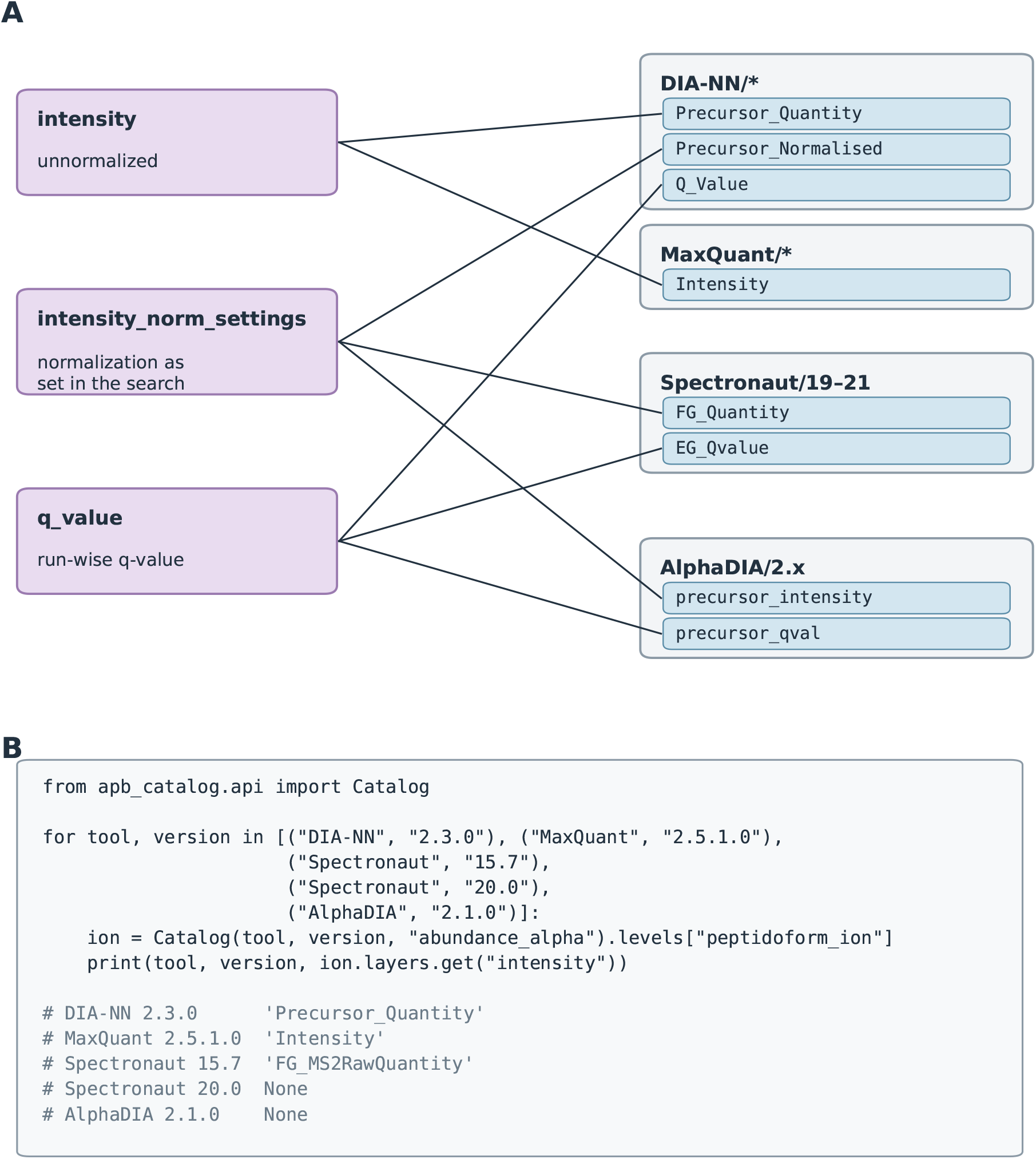
The catalog maps one term to a different field in each quantification tool. (A) Three terms of the abundance and confidence sets and the peptidoform-ion field that each maps to in `DIA-NN`, `MaxQuant`, `Spectronaut` 19–21, and `AlphaDIA` 2.x. Neither `Spectronaut` 19–21 nor `AlphaDIA` has a field for unnormalized intensity. (B) An application asks for the term `intensity` and gets the field of each tool version, or `None`. `Spectronaut` reports an unnormalized intensity only in version 15. `apb-catalog` computed the output lines. Section S9 describes the columns of the catalog and how its tables relate.

The catalog viewer (https://fgcz-ms.uzh.ch/public/apb-catalog/) runs in the browser and shows, for each quantification tool and version, the column that each term maps to and the terms that its results lack. Its editor, a demonstration, lets a user map a term to a field for a quantification tool and version, change such a mapping, or start a new set; it checks the edited tables against the catalog’s table schema, accepts only tool versions that a parsing rule declares, and saves the tables as CSV files.

A consumer is any program that reads a ProteoBridge result; some add what they compute to it, such as FASTA matches or benchmark scores, and new consumers need no change to ProteoBridge. `apb-fasta` matches peptides against any FASTA files, typically those of the search, through the `ProForma peptide` columns, and uses `fasta accessions` and `protein assignment` to compare the reported proteins and protein groups with these matches. It reads species and contaminants from the FASTA headers, the species from the UniProtKB `OS=` field or the bracketed organism of RefSeq headers, and contaminants and decoys from accession prefixes such as `CON`, `Cont` , `REV` , or `DECOY` , and records species, contaminant, and decoy matches in `varm[“fasta validation”]`. `apb-proteobench` scores a ProteoBridge result as `ProteoBench` does; it supports nine quantitative modules, and the corpus covers seven (Table S3). The user names the module, such as `dia singlecell` for low-input DIA of human and yeast proteins, and the module’s sample table, which records the quantity of each species in each sample, sets the expected ratios. On request, it adds the module’s sample design and verifies the peptides with `apb-fasta`; it then computes from the primary abundance the benchmark metrics that `ProteoBench` reports, such as the error of the spike-in fold changes against their known ratios (Section S10).

`apb-export` generates application-specific input files using export rules (Section S11.1). These rules reference roles, ProteoBridge columns (for example, ProForma identifiers), and catalog terms, rather than the column names of a quantification tool. As an example, we export a `DIA-NN` report that was searched against the `ProteoBench` FASTA as input for `prolfquapp` (Figure 5):

**Figure 5:**
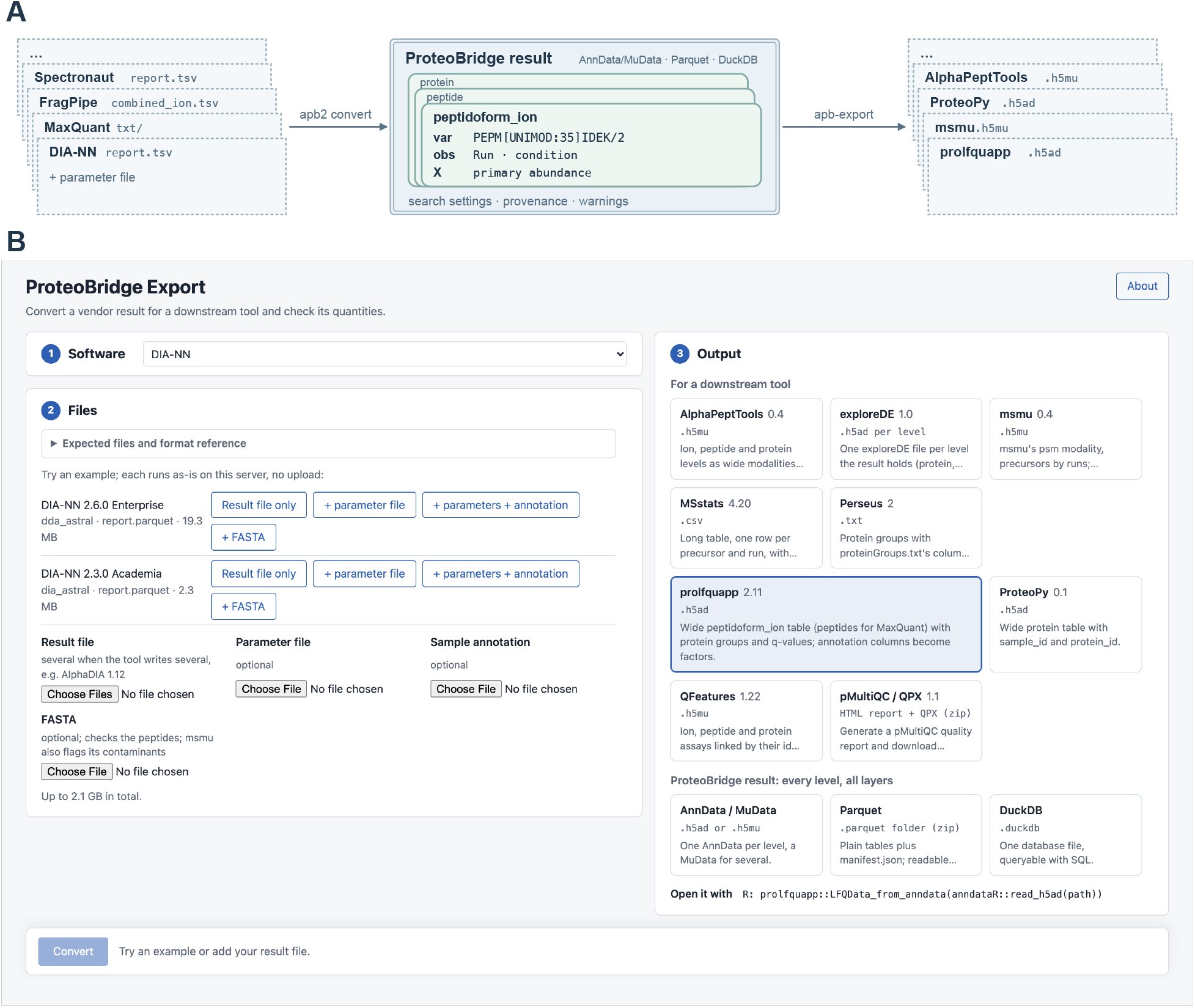
(A) Route of the export example: software output passes through `apb2 convert` into the ProteoBridge result, and `apb-export` writes the input of applications. (B) The ProteoBridge export web app with `DIA-NN` selected and `prolfquapp` as output.

`apb-export prolfqua report.tsv prolfqua.h5ad \`

`--params report.log.txt \`

`--annotation dia_aif.sdrf.tsv \`

`--fasta ProteoBenchFASTA_MixedSpecies_HYE.fasta`

The command converts the report and adds the sample table; from the FASTA file, it adds the protein annotations that `prolfquapp` needs, such as protein descriptions and lengths, and exports an AnnData file for `prolfquapp`. The same steps also run one at a time from Python (Section S11.2), and Section S11.3 lists the catalog choices of the export rule.

Besides `prolfquapp`, ProteoBridge currently writes nine other export targets, among them `msmu`, `ProteoPy`, and `AlphaPeptTools` (see Section S11.1). On a synthetic `DIA-NN` report, `prolfquapp` and `msmu` each compute the same peptide and protein values from their ProteoBridge export as from their own `DIA-NN` reader (Section S11.4). The ProteoBridge export web app runs the same exports without installation (Figure 5B; Section S12).

We are working on support for reports of post-translational modification (PTM) sites. Several quantification tools, including `Spectronaut`, `FragPipe`, and `DIA-NN`, write separate site reports, including localization probability and often the sequence window for each modification site. A site level in ProteoBridge would provide PTM analyses, such as integrated analysis of modification sites and the total proteome^66^, with one shared reader. Sites relate to peptidoforms many to many: a peptidoform can carry several sites, and a site can lie on several peptidoforms. `mulink`, which stores links between MuData modalities, could hold this relation^67^.

## Conclusions

ProteoBridge converts the result tables of every quantification tool into annotated data matrices, one per level, but keeps each parsed column under a name taken from the tool’s header. A user who knows a tool’s output layout thus finds all of its data under names close to the original headers. Applications then read all tools through one API. Because ProteoBridge stores results as HDF5, Parquet, or DuckDB, which most languages read, the API has ports to R and TypeScript. `mzAPI` left open how one API exposes what only one tool writes. ProteoBridge answers with a catalog, which maps a term, such as unnormalized ion intensity, to each tool’s column. An application chooses the sets of the catalog that it uses and can keep its interface minimal: it reads every quantification tool whose results hold the terms it requires. `apb-export` currently uses the catalog to write ten export targets. A new quantification tool or version needs a parsing rule and catalog entries, and a new term only catalog entries. Neither changes the parser or the consumers. The parsing rules are tested on public inputs. Together with the catalog, they are openly offered as the basis of a community resource for reading the outputs of quantification software.

## Software and data availability

The packages are open source in the anndata-omics-bridge GitHub organization and require Python 3.13 or later. `apb2`, `apb-fasta`, `apb-proteobench`, and `apb-catalog` are available from PyPI (pypi.org/project/apb2, apb-fasta, apb-proteobench, apb-catalog) and are archived on Zenodo^36,68–70^. `apb-export` and the export web app `apb-export-ui` (Section S12) are also available from PyPI (apb-export, apb-export-ui). `APB Studio` and the R and TypeScript ports of `apb-catalog`, `apbCatalogR` and `apb-catalog-ts`, are available from the organization’s repositories. The ProteoBridge export web app (https://apps-dev.bfabric.org/apb-export/) converts uploaded software output to any export target or storage format, the `APB Studio` viewer (https://fgcz-ms.uzh.ch/public/apb-studio/) shows the saved corpus runs, the catalog viewer (https://fgcz-ms.uzh.ch/public/apb-catalog/) shows the catalog, and the parsing-rules viewer (https://fgcz-ms.uzh.ch/public/apb-rules/) shows the parsing rules. The documentation (https://anndata-omics-bridge.github.io/apb2/) describes installation, supported inputs, and the Python interface. The `ProteoBench` result repositories on GitHub (github.com/Proteobench) index the submissions, and we downloaded them from the ProteoBench server^10^. The `directLFQ` exports are available from their Zenodo mirror^71^. The `Spectronaut` and `DIA-NN` reports analyzed in the `MSstats+` publication^42^ are available from a second Zenodo mirror^72^. Their raw data are found on ProteomeXchange and PRIDE^73,74^.

## Author contributions

R.D., R.B., S.v.P., H.W., W.E.W., C.J., and M.L.-P. developed the `ProteoBench` parameter and result parsing on which ProteoBridge is built. R.D., R.B., C.J., and W.E.W. developed the `ProteoBench` quantification benchmark modules. L.S. contributed to the initial design and conceptualization of ProteoBridge. C.P., H.R., and L.M. supervised the work. W.E.W. conceived and developed ProteoBridge and performed the analyses. W.E.W., R.D., and M.L.-P. wrote the manuscript. All authors reviewed the manuscript.

## Acknowledgments

M.L.-P. was supported by grants from the French National Research Agency (ProFI projects: ANR-10-INBS-08 & ANR-24-INBS-0015).

S.v.P. acknowledges funding from the Research Foundation Flanders (FWO) [1SA7M26N].

R.D. and R.B. acknowledge funding from the Research Foundation Flanders (FWO) [1SH9O24N, 12A6L24N]. C.J. acknowledges funding from the European Union’s Horizon Europe under grant agreement no. 101119980.

We thank the `ProteoBench` contributors for the parameter, result, and modification parsing code, and the submitters of the public workflow outputs used for testing. We thank Lucas Diedrich (`AlphaPeptTools`), Georg Wallmann (`AlphaDIA`), Oliver Bernhardt (`Spectronaut`), and Sunghyun Huh (`msmu`) for checking how we describe their software, Isabell Bludau for reviewing the manuscript, and Isabell Bludau and Ian Fichtner for testing the `ProteoPy` export. We thank Mikhail A. Bakunin (1814–1876), who in the matter of boots deferred to the authority of the bootmaker; ProteoBridge leaves what a column means to the developer of each quantification tool, and what an application needs to its developer.

## Notes

The authors declare no competing financial interest.

## AI use disclosure

We used the coding assistants OpenAI `Codex` and Anthropic `Claude Code`, and assistants built on `Mistral` and `Qwen` models. They assisted with software design and implementation, figure generation, and the drafting of this manuscript. These tools do not meet authorship criteria, and we do not list them as authors. We directed all AI use and reviewed the resulting code and text. The authors take full responsibility for the content of this article.

## Supporting Information

The *Supporting Information* explains the bootmaker of Figure 1 (S1), gives a glossary (S2), and compares converters (S3). It lists the inputs and their accessions (S4), reports performance and storage measurements (S5), and describes quality checks and corpus runs (S6). It lists the quantification tools with their citations, preparation steps, and further notation examples (S7), and the settings that the parameter parsers recovered (S8). It describes the catalog with code examples (S9) and gives the `ProteoBench` scoring example (S10). It closes with the export rules, the export example, and the export checks (S11), and the export web app (S12).

## For Table of Contents Only

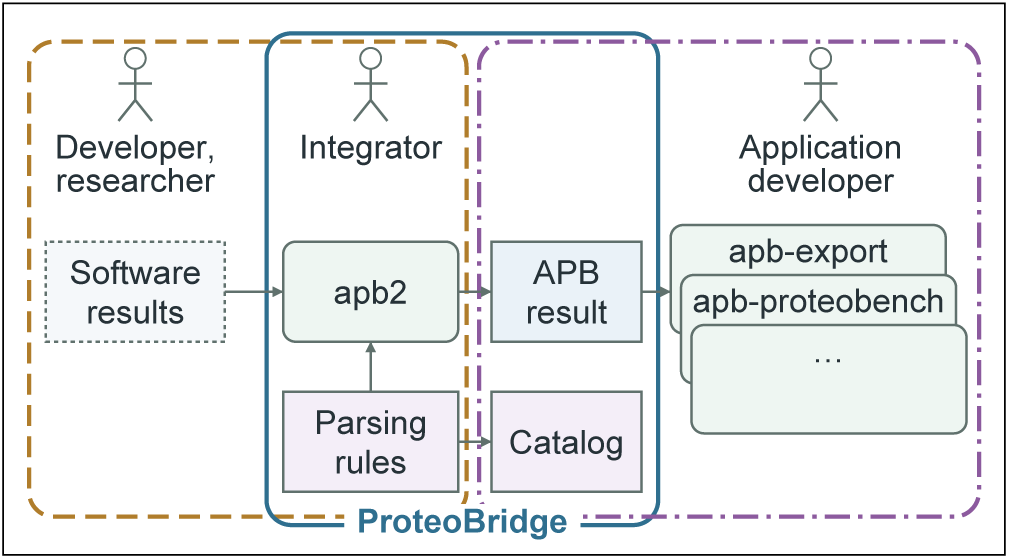

