## Supporting Information for "ProteoBridge Connects Proteomics Quantification Outputs to Reusable Analysis Workflows"

#*Univ Toulouse, CNRS, IPBS, Toulouse, France*

@*Infrastructure Nationale de Protéomique, ProFI, UAR 2048, Toulouse, France*

**Abstract**

This Supporting Information gives a glossary and compares converters. It lists the inputs with their raw-data accessions, reports the conversion and storage measurements, and describes the quality checks and corpus runs. It describes the parsing rules, with preparation steps and a link to the parsing-rules viewer, and the search settings that the parameter parsers recovered. It shows the abundance and identification-confidence sets of the catalog with code examples, and gives the ProteoBench scoring example. It closes with the export rules, the export example in Python, the export checks, and the export web app.

### S1 The bootmaker

Main-text Figure 1 and the Acknowledgments cite Mikhail Bakunin's bootmaker. In *God and the State*, written in 1871 and published in 1882, Bakunin asks whether he rejects all authority and answers: "In the matter of boots, I refer to the authority of the bootmaker; concerning houses, canals, or railroads, I consult that of the architect or engineer." He accepts this authority because his own reason grants it, and he consults several specialists, compares their opinions and keeps the right to criticize them: "But I allow neither the bootmaker nor the architect nor the savant to impose his authority upon me." [1]

ProteoBridge divides expertise in the same way. The quantification software developer knows what a column of the tool means, and the application developer knows what the application needs. The integrator connects the two through parsing rules and catalog entries and decides neither (main-text Figure 1).

Bakunin also rejected rule by experts, a body of specialists that decides for everyone: "A scientific body to which had been confided the government of society would soon end by devoting itself no longer to science at all, but to quite another affair." [1] In the Paris Commune of 1871 he saw order built "from the bottom up", by a federation of self-governing communes in place of a central state[2].

ProteoBridge keeps each tool's columns under names taken from the tool's headers. The catalog maps terms to them, and its sets, the parsing rules and the export rules are open files that anyone is invited to extend; for the catalog, the editor of the catalog viewer shows how to add a mapping or a set (main-text Results). A standard defined by PSI would become another export target, written by its own export rule, for the applications that need it. As `mzAPI` proposed common access to raw data instead of a common file[3], ProteoBridge offers common access to quantification results.

#### S2 Glossary

Table S1 defines the terms used throughout the manuscript and figures.

Table S1: Terms of the article.

| Term | Definition |
|---|---|
| Actor | Person with an interest in ProteoBridge: quantification software developer, integrator, or application developer |
| Annotated data matrix | The data model of one level: a matrix of quantities with a feature table that annotates its rows and a sample table that annotates its columns, as in ExpressionSet, SummarizedExperiment, or AnnData; MuData and MultiAssayExperiment group several |
| Application | Software that analyzes or uses results, such as `prolfquapp`, `msmu`, or `ProteoBench` |
| Application developer | Builds one application. States the terms it needs in the catalog and writes its export rule against roles, ProteoBridge's own columns, and catalog terms; never sees the output layout of a quantification tool |
| Catalog | The terms of `apb-catalog`, kept in sets, with the layer or column that each term maps to for each quantification tool and version |
| Consumer | Software that reads a ProteoBridge result, such as `apb-fasta`, `apb-proteobench`, and `apb-export` |
| Data model | The structure of a result: quantity matrices with tables that annotate their rows and columns, as in AnnData, MuData, or SummarizedExperiment |
| Export rule | JSON file that maps the ProteoBridge result to the input of one application, through roles, ProteoBridge's own columns, and catalog terms; from the parsing rule recorded in the result it reads only the software name, to choose the level |
| Export target | The file that an export rule writes for one application |
| Field | Layer or column of the ProteoBridge result; the catalog returns its name, such as `Precursor_Quantity` |
| File format | How data are stored as bytes, without domain meaning, such as XML, TSV, Parquet, HDF5, or DuckDB; storage format for the three that ProteoBridge writes |

| Term | Definition |
|---|---|
| Integrator | Connects many quantification tools to several downstream applications. Knows what applications need but not every application, and hands them the catalog and exports. Writes the parsing rules and parameter parsers and fills the catalog entries for each tool and version; each new tool needs its own rules and entries, and an application may ask for new terms |
| JSON summary | File beside every ProteoBridge result that lists its levels, columns, layers, roles, provenance records, and warnings; a JSON Schema defines it |
| Key | Name of a term, such as `intensity_norm_settings`; it fixes the term's kind and qualifiers |
| Layer | One quantity of a level, such as an intensity or a q-value, as a matrix of samples by features |
| Level | One feature type of the ProteoBridge result, such as peptidoform ions or proteins: an annotated data matrix with its own sample table, feature table, and layers |
| Output layout | The tables, columns, and notation that one quantification tool and version writes, such as a `DIA-NN` report |
| Parameter parser | Python code that reads the settings of one quantification tool into one typed record |
| Parsing rule | JSON file that describes one output layout of one quantification tool: it selects the identifier columns and quantities and declares the roles, the modification map, and the missing-value and duplicate policies |
| ProteoBridge result | The converted result: one level per feature type, each a sample table, a feature table, and aligned quantity layers, with the provenance of every step |
| Quantification software developer | Builds a bottom-up MS quantification tool and needs to know nothing about ProteoBridge |
| Quantification software, quantification tool | Software whose result tables and parameter files ProteoBridge reads, such as `DIA-NN` or `MaxQuant` |
| Resolver | Code in `apb-catalog` that answers a consumer's request for a term with the layer or feature-table column it maps to, with "none", or by stopping when the answer is ambiguous or unknown |
| Role | A mark that a parsing rule declares on a column or layer that every quantification tool reports in the same sense: `protein_assignment`, `fasta_accessions`, or `abundance`; `abundance` marks every abundance layer, intensities and iBAQ alike |
| Sample table | The table of samples and their experimental design that the researcher supplies, as SDRF-Proteomics or a `prolfquapp` table; also the sample table (`obs`) of each level of the ProteoBridge result, to which `apb2 annotate` adds that design |
| Set | The terms of one topic in the catalog, such as abundance; several consumers use one set |
| Shared parser | Code in `apb2` that applies every parsing rule in the same steps and order |
| Standard | Agreement on what the columns or elements of a file mean, such as mzML, mzTab, mzPeak, QPX, or SDRF-Proteomics; ProteoBridge defines none |
| Term | What an application asks the catalog for, by its key, such as unnormalized ion intensity; the catalog maps it to a field for each quantification tool and version |
| Warning | Problem that a step records in the result instead of stopping or silently dropping data |

### S3 Converters compared

Table S2 compares ProteoBridge with four converters from quantification software output to a common data model or standard, as of 27 October 2026: QPX 1.1.6, and `AlphaPeptTools`, `msmu`, and `ProteoPy` as in their repositories in September and October 2026. Tools read counts the quantification tools that each converter reads; `ProteoPy` reads `DIA-NN`, and any other long table through a column map that the user writes. Levels are the feature types of the result: all, every level the tool reports; subset, some of the levels the tool reports; derived, coarser levels that the converter computes, as `msmu` computes its protein level and `ProteoPy` sums `DIA-NN` precursors to peptidoforms, peptides, and proteins. Feature index is the identifier of each feature: ProForma, converted from each tool's notation; tool ID, the tool's own identifier, such as `DIA-NN`'s `Precursor.Id`. Quantities are the layers of a level: all, every value per run and feature; primary + 2, one chosen intensity and at most two further abundances, fixed per tool, such as MaxLFQ or iBAQ; one, a single quantity per AnnData object, which `AlphaPeptTools` lets the user choose. Feature information: all, every column that carries information; fixed, a set schema; fixed + user-defined, a set schema plus the report columns that the user names when calling the converter; fixed + report, a set schema plus the raw report without stated meanings. Level links, how a feature points to its features at a coarser level: keys, key columns in each row; matrix, relation matrices between levels; inferred, a peptide-to-protein map from protein inference. Parameters: parsed, the parameter file read into one typed record; not read, the parameter file left out. Output: several, AnnData/MuData, Parquet, DuckDB, and the `apb-export` export targets.

For raw MS data, `MsBackendRawFileReader` implements the `MsBackend` interface of `Spectra` for Thermo files through `rawrr`[4,5]; it exposes a common set of spectrum variables, a subset of what the raw file records.

Table S2: Converters from quantification software output to a common data model or standard, as of 27 October 2026.

| | APB | QPX | AlphaPeptTools | msmu | ProteoPy |
|---|---|---|---|---|---|
| Tools read | 14 | 8 | 8 | 5 | 1 |
| Levels | all | subset | subset | derived | derived |
| Feature index | ProForma | ProForma | tool ID | tool ID | tool ID |
| Quantities (layers) | all | primary + 2 | one | one | one |
| Feature information | all | fixed | fixed + user-defined | fixed + report | fixed |
| Level links | keys + matrix | keys + matrix | matrix | inferred | keys |
| Parameters | parsed | not read | not read | not read | not read |
| Output | several | Parquet + MuData | AnnData/MuData | AnnData/MuData | AnnData/MuData |

### S4 Inputs and accessions

Main-text Figure 3A,B and Section S5.2 rest on 203 inputs: 198 `ProteoBench` quantification submissions and five large exports from the `directLFQ` and `MSstats+` benchmarks (Table S3). Main-text Figure 3C and Section S5.3 rest on 19 inputs: 12 `ProteoBench` submissions, the `MaxQuant` export of PXD070049, the `MetaMorpheus` IonStar export, and the five large inputs. Three of them, a `Sage` submission, the `MaxQuant` export, and the IonStar export, are not among the 203.

Table S3: Conversion inputs by `ProteoBench` module and quantification tool; rows outside the `ProteoBench` modules are the large inputs. Input size is the stored size, gzip-compressed for text inputs; input size and ion count are medians per row.

| Module | Quantification tool | Inputs | Median input (MB) | Median ions |
|---|---|---|---|---|
| dda_astral | DIA-NN | 2 | 58.8 | 67418 |
| dda_astral | FragPipe | 2 | 6.1 | 52305 |
| dda_astral | MaxQuant | 1 | 12.6 | 24987 |
| dda_astral | i2MassChroQ | 3 | 5.8 | 37348 |
| dda_qexactive | AlphaPept | 4 | 59.2 | 58718 |
| dda_qexactive | FragPipe | 7 | 6.9 | 83007 |
| dda_qexactive | MS-Angel | 1 | 40.6 | 67048 |
| dda_qexactive | MaxQuant | 15 | 23.2 | 53588 |
| dda_qexactive | PEAKS | 1 | 10.0 | 76129 |
| dda_qexactive | Proline | 1 | 23.0 | 60499 |
| dda_qexactive | WOMBAT-P | 5 | 2.4 | 48729 |
| dda_qexactive | i2MassChroQ | 10 | 10.2 | 73828 |
| dda_qexactive | quantms | 1 | 2.6 | 48794 |
| dia_aif | DIA-NN | 8 | 92.6 | 97974 |
| dia_aif | FragPipe (DIA-NN quant) | 5 | 121.6 | 74692 |
| dia_aif | PEAKS | 1 | 14.4 | 112140 |
| dia_aif | Spectronaut | 3 | 122.3 | 103747 |
| dia_astral | AlphaDIA | 16 | 55.6 | 119658 |

| Module | Quantification tool | Inputs | Median input (MB) | Median ions |
|---|---|---|---|---|
| dia_astral | DIA-NN | 32 | 73.3 | 127786 |
| dia_astral | FragPipe (DIA-NN quant) | 5 | 154.1 | 91287 |
| dia_astral | MaxQuant | 1 | 13.4 | 24990 |
| dia_astral | PEAKS | 8 | 12.7 | 146457 |
| dia_astral | Spectronaut | 8 | 153.9 | 120826 |
| dia_diapasef | AlphaDIA | 7 | 56.6 | 89282 |
| dia_diapasef | DIA-NN | 15 | 95.2 | 120283 |
| dia_diapasef | FragPipe (DIA-NN quant) | 7 | 166.3 | 86865 |
| dia_diapasef | PEAKS | 6 | 12.0 | 114934 |
| dia_diapasef | Spectronaut | 7 | 171.8 | 131920 |
| dia_singlecell | AlphaDIA | 1 | 10.2 | 28122 |
| dia_singlecell | DIA-NN | 6 | 23.0 | 49310 |
| dia_singlecell | FragPipe (DIA-NN quant) | 3 | 56.1 | 32036 |
| dia_singlecell | MaxQuant | 2 | 14.8 | 29753 |
| dia_zenotof | AlphaDIA | 2 | 73.9 | 65604 |
| dia_zenotof | DIA-NN | 1 | 55.7 | 104781 |
| dia_zenotof | PEAKS | 1 | 15.1 | 136142 |
| directlfq | MaxQuant | 1 | 1128.8 | 59515 |
| directlfq | Spectronaut | 1 | 1338.9 | 181147 |
| msstats_plus | DIA-NN | 2 | 5918.1 | 308185 |
| msstats_plus | Spectronaut | 1 | 4816.1 | 286415 |

Every input traces back to raw data in a public PRIDE project (Table S4). We checked each accession through the PRIDE and ProteomeXchange interfaces and found the run names of every input in the project's file list or SDRF. ProteoBench names no accession for the DDA Astral module yet; its six run files match PXD070049 by name and byte size. The Spectronaut LargeFC report derives from the raw files of Bruderer et al.[6], deposited as PXD005573.

Table S4: Raw-data accessions of all inputs, with the number of ProteoBench submissions per module. HYE: human, yeast, and *E. coli* mixture (A 65:30:5, B 65:15:20).

| Input | Instrument and acquisition | Accession | Public since |
|---|---|---|---|
| ProteoBench dda_qexactive (45) | Q Exactive HF-X; DDA; HYE | PXD028735 | 2022-02-14 |
| ProteoBench dda_astral (8) | Orbitrap Astral; DDA, 15 min, 50 ng; HYE | PXD070049 | 2026-02-03 |
| ProteoBench dia_aif (17) | Q Exactive HF-X; DIA, all-ion fragmentation; HYE | PXD028735 | 2022-02-14 |
| ProteoBench dia_astral (70) | Orbitrap Astral; DIA, 15 min, 50 ng; HYE | PXD070049 | 2026-02-03 |
| ProteoBench dia_diapasef (42) | timsTOF SCP; diaPASEF, 25 ng; HYE | PXD062685 | 2025-05-09 |
| ProteoBench dia_singlecell (12) | Orbitrap Astral; DIA with FAIMS, low input; HeLa and yeast | PXD049412 | 2025-01-16 |

| Input | Instrument and acquisition | Accession | Public since |
|---|---|---|---|
| ProteoBench dia_zenotof (4) | ZenoTOF 8600; Zeno SWATH, 15 min, 50 ng; HYE | PXD070049 | 2026-02-03 |
| MetaMorpheus IonStar export | Orbitrap Fusion; DDA, HCD; human with E. coli spike-in at five levels | PXD003881 | 2018-05-16 |
| MaxQuant export | Orbitrap Astral; DIA, 15 min, 50 ng; human digest | PXD070049 | 2026-02-03 |
| Spectronaut LargeFC report | Q Exactive HF; DIA, 240 min; four-species mixture | PXD005573 | 2017-10-27 |
| MaxQuant 200 HeLa replicates | Q Exactive HF-X; DDA; HeLa digest | PXD015087 | 2019-12-09 |
| DIA-NN K562 dilution series | Orbitrap Astral; DIA; K562 digest dilutions | PXD066486 | 2025-10-01 |
| Spectronaut mixture of proteomes | Orbitrap Astral; narrow-window DIA; HYE in six ratios | PXD046444 | 2024-02-01 |
| DIA-NN Pan-Cancer Proteome Atlas | Q Exactive HF; DIA; tumor tissues | PXD054790 | 2025-05-30 |

The directLFQ authors distribute their benchmark inputs through download links without a DOI[7], and two MSstats+ reports sit only inside archives[8]. We deposited gzip-compressed copies with checksums on Zenodo[9,10] and converted them without parameter files. The TPCPA report is the native DIA-NN 1.8 report from PXD054790.

### S5 Performance and storage measurements

#### S5.1 Computer and measurement

We measured conversion on 21 November 2026 on an Apple M4 Pro with 48 GiB RAM under macOS 27.2.1. Input size is the stored size of the result files that the parsing rule reads, without parameter or FASTA files. Text inputs are stored gzip-compressed, so their conversion time includes decompression. Peak memory is given in GB ($10^9$ bytes), rounded to 0.1 GB. apb2 0.1.3 ran with Polars 1.44.1 and four threads, one conversion at a time. We measured each input once per route and did not control filesystem caches.

#### S5.2 Conversion time and memory

Main-text Figure 3A,B shows the conversions to all reported levels. Figure S1 shows both routes for the same 203 inputs against ion count.

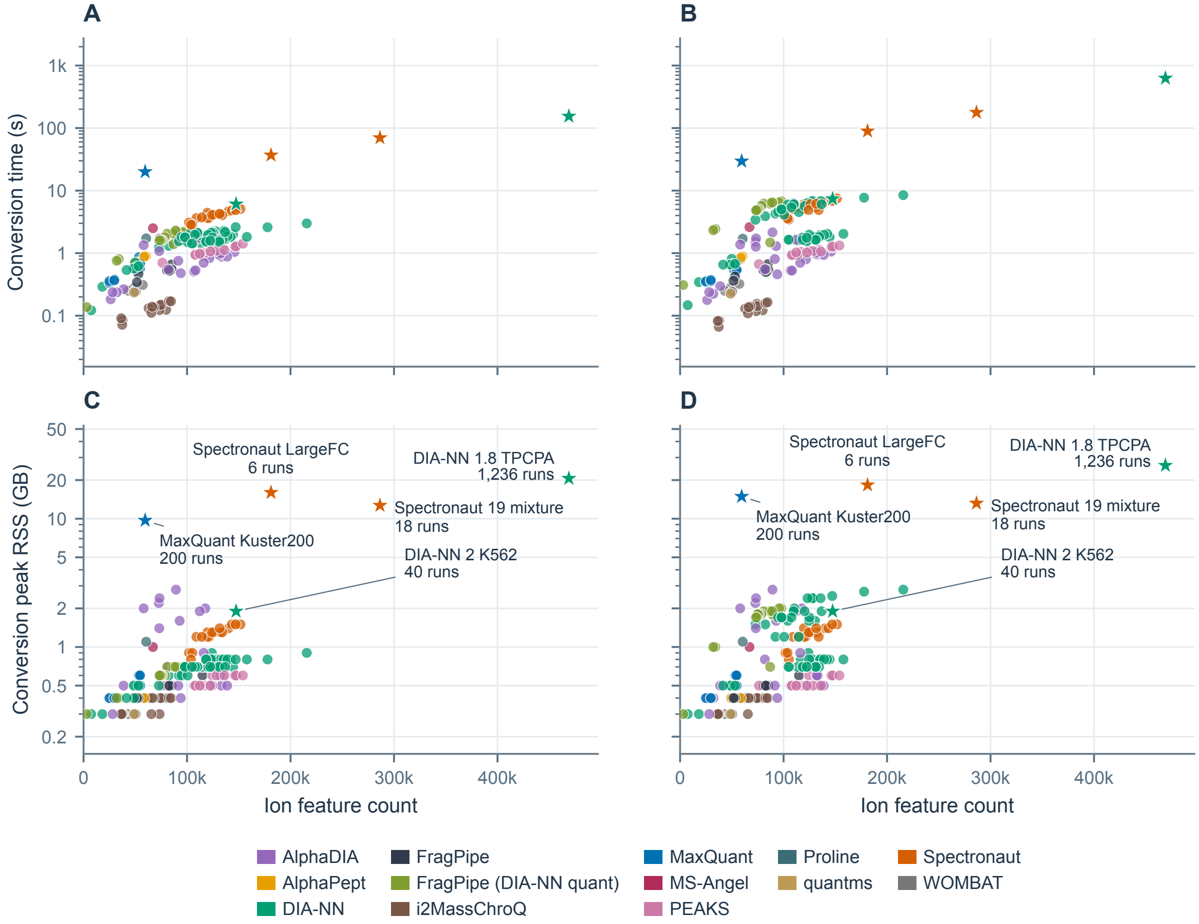


Figure S1: ProteoBridge conversion versus ion count for the 203 inputs of main-text Figure 3: 198 ProteoBench submissions with six samples each (circles) and five large inputs (stars, labeled with their run counts in C and D). (A,C) Ion-only conversion time and peak process RSS. (B,D) Conversion to all reported levels, written as MuData: time and peak process RSS. Color identifies the quantification tool, as in main-text Figure 3. Ion-count axes are linear; time and RSS axes are logarithmic.

#### S5.3 Storage sizes and read-back

Figure S2 shows the absolute input and output sizes of the storage selection. The measured apb2 source compresses the AnnData/MuData arrays with gzip. Against the input, the summed sizes are 91.4% for AnnData/MuData, 86.0% for Parquet, and 159.3% for DuckDB, and the medians of the per-dataset ratios are 103.3%, 64.7%, and 87.6%; the sums weight each input by its size, the medians weight every input equally.

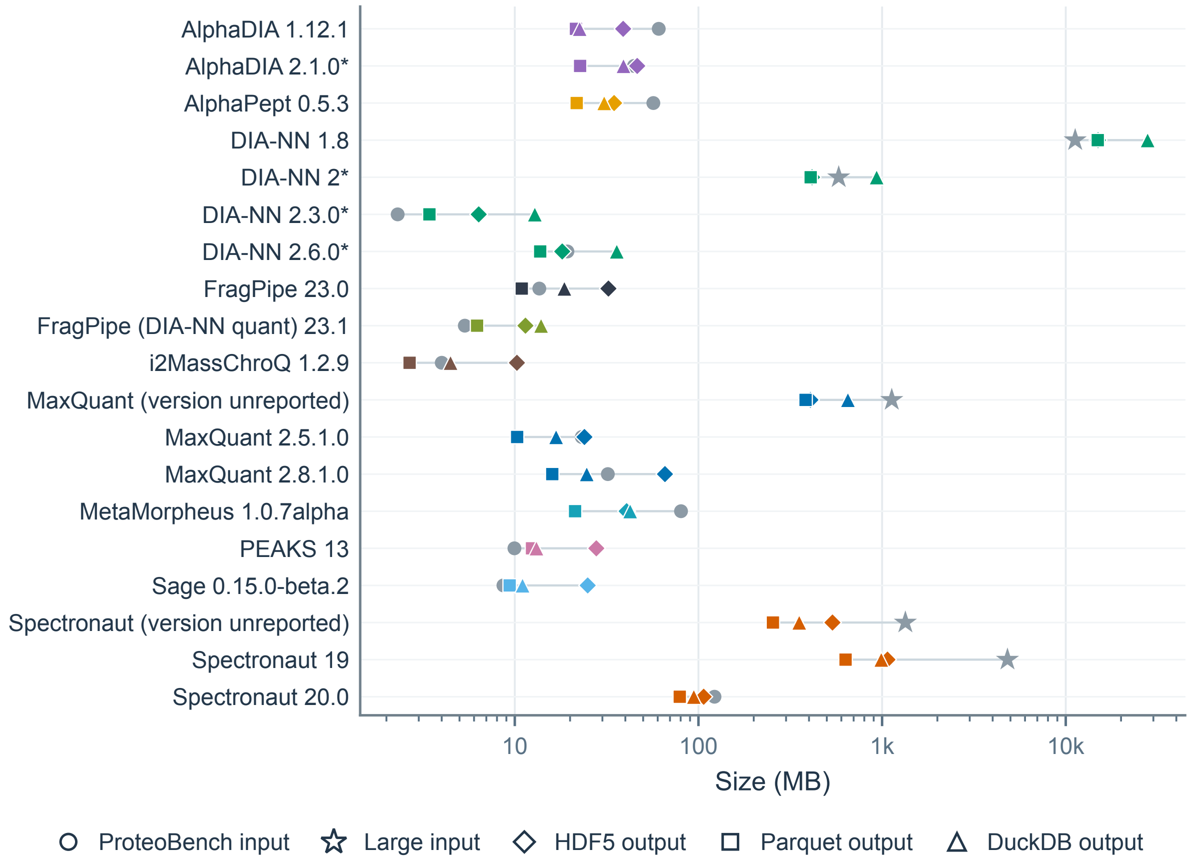


Figure S2: Absolute input and output sizes for the storage datasets of main-text Figure 3C, in decimal megabytes on a logarithmic axis: AnnData/MuData in HDF5 (diamonds), Parquet (squares) and DuckDB (triangles). Marker color identifies the quantification tool. Gray circles mark `ProteoBench` inputs and gray stars the large inputs; thin lines join the sizes of one dataset. An asterisk marks an input that is already a Parquet file.

Reading the results back through the APB Python interface gives identical level names, dimensions, and identifiers for all three formats. All 547 annotation columns and all 295 quantity layers, about 20.9 billion cells, agree in values and missing-value positions. As AnnData specifies, HDF5 stores missing floating-point cells as NaN, not null.

### S6 Quality checks and corpus runs

Table S5 lists the checks of each APB package and of `protein-fasta` and `prozor`, the two libraries that `apb-fasta` builds on. The APB packages share one `Ruff`[11] rule selection, except `APB Studio`, whose own selection adds limits on arguments, branches, and statements. On every push, GitHub continuous integration runs `make check` in each package, and the pre-commit and pre-push hooks in `APB Studio`. Each package exposes its interface in one `api.py` module, the only module that other packages import; `import-linter` contracts and tests enforce this boundary.

Table S5: Quality checks per APB package and for `protein-fasta` and `prozor`; APB package columns omit the `apb-` prefix. Numbers are `import-linter` contracts, test functions, or branch-coverage floors, as of 11 November 2026; `twine` checks the built distributions.

| Check | apb2 | fasta | proteobench | catalog | export | Studio | protein-fasta | prozor |
|---|---|---|---|---|---|---|---|---|
| `Ruff` format and lint | yes | yes | yes | yes | yes | yes | yes | yes |
| `Pyright` strict | yes | yes | yes | yes | yes | yes; type-ignore comments rejected | yes | yes |
| `deptry` dependency check | yes | yes | yes | yes | yes | yes | yes | yes |
| `import-linter` contracts | 14 | 2 | 3 | none | none | none | 11 | 2 |
| Sibling imports only through `api.py` | no siblings | test | test | test | test | test | no siblings | no siblings |
| Test functions | 910 | 46 | 91 | 180 | 118 | 204 | 306 | 33 |
| Branch-coverage floor | reported | 80% | 80% | 80% | 80% | 90%; diff coverage | 80% | 80% |
| Strict docs build | yes | yes | yes | yes | yes | yes | yes | yes |
| Package build check | `twine` | `twine` | `twine` | `twine` | `twine` | smoke test | `twine` | `twine` |

The `APB Studio` viewer (`https://fgcz-ms.uzh.ch/public/apb-studio/`) shows the corpus runs behind main-text Figure 3, the `pmultiqc` reports of the 198 `ProteoBench` submissions, and a corpus of inputs that fail, or convert with warnings, for a known reason. Its runs use a newer `apb2` than the measured source, so their times differ from those of main-text Figure 3. Running all corpus workflows under coverage on 6 October 2026 executed 83% of apb2's 7,473 statements; we removed about 30 statements that had no caller. Besides the conversion warnings, FASTA matching records peptides and protein-group members absent from the FASTA, and ambiguous members; annotation records sample names that it matched only approximately. Corpus runs also led to rule fixes. The DIA-NN rules did not map the phosphorylation and GlyGly notations UniMod:21 and UniMod:121. The PEAKS cells - and the WOMBAT-P cells NA were unreadable until the rules declared them missing. The i2MassChroQ rule read peptides from a column with PSI-MOD annotations, which left 16–24% of them unmatched.

### S7 Parsing rules

#### S7.1 Quantification tools and rule generations

Parsing rules currently cover MaxQuant, DIA-NN, and FragPipe[12–16], Spectronaut[17], PEAKS[18], MetaMorpheus[19], i2MassChroQ[20], Proline[21], MS-Angel[22], quantms[23], WOMBAT-P[24], AlphaDIA, AlphaPept, and Sage[25–27]. The apb2 documentation lists the supported versions, file layouts, and levels of each. Spectronaut 20 and 21 can also write their report as Parquet; the Spectronaut parsing rules currently read the TSV report. The parsing-rules viewer (https://fgcz-ms.uzh.ch/public/apb-rules/) shows each packaged parsing rule level by level: the columns it selects for the sample table, the feature table, and the quantity layers, and the roles, the modification map, and the missing-value and duplicate policies it declares.

One quantification tool can need several parsing rules. DIA-NN currently has three. Up to version 1.7, it writes a TSV report without posterior error probabilities (PEP), library protein-group q-values (Lib.PG.Q.Value), and PTM columns, which versions 1.8 and 1.9 add. Version 2 writes only Parquet, adds top-N quantities (PG.TopN, Genes.TopN) and drops the normalized gene and protein quantities (PG.Normalised,

Genes.Normalised). The software version in the parameter file selects the rule. Without a parameter file, the report columns select it: 1.7 reports lack Lib.PG.Q.Value, 1.8 and 1.9 reports have PG.Normalised, and version 2 reports have PG.TopN.

##### S7.2 Preparation steps

Most parsing rules describe one table. A few output layouts need a preparation step, a short Python function that runs before the rule. For MaxQuant, it joins the modification-specific peptide, peptide, and protein-group tables, and gives each evidence row the protein group of its leading razor protein. For AlphaDIA 1.12, it joins the intensities (precursor.matrix.tsv) with the precursor metadata (precursors.tsv). For MS-Angel, it joins the peptide-ion sheet of a workbook with its protein-set sheet. MetaMorpheus writes FlashLFQ peaks (AllQuantifiedPeaks.tsv), one row per MS1 peak and run. Its step drops peaks that FlashLFQ assigns to several peptides, and match-between-runs peaks taken at a random retention time. The WOMBAT-P step drops the same multi-peptide rows. The Proline Studio export needs no preparation step: its parsing rule joins the sameset and subset accessions of each row into one Proteins column, which carries the fasta_accessions role.

##### S7.3 Notations and parameter files

Figure S3 shows how the tools write three peptidoforms and four settings.

#### S8 Parameter parsers and recovered settings

Besides the settings named in the main text, the typed record[28] of settings holds the FDR thresholds, maximum charge, and minimum peptide length. Figure S4 shows which settings the parsers recovered from the 198 ProteoBench submissions as of 21 November 2026. Enzyme, missed cleavages, modifications, PSM FDR, and both mass tolerances were recov-

## A

##### As written by the software

| | Oxidized methionine<br>variable modification | Carbamidomethylated cysteine<br>fixed modification | Protein N-terminal acetylation<br>terminal modification |
|---|---|---|---|
| DIA-NN | AM(UniMod:35)DVYQK | ALIQC(UniMod:4)AK | (UniMod:1)MEAFQELR |
| AlphaDIA | AMDVYQK; Oxidation@M; 2 | ALIQCAK; Carbamidomethyl@C; 5 | MEAFQELR; Acetyl@Protein_N-term; 0 |
| FragPipe (DIA-NN quant) | AM(UniMod:35)DVYQK | ALIQC(UniMod:4)AK | (UniMod:1)MEAFQELR |
| MaxQuant | _AM(Oxidation (M))DVYQK_ | _ALIQCAK_ fixed modification not written | _(Acetyl (Protein N-term))MEAFQELR_ |
| Spectronaut | _AM[Oxidation (M)]DVYQK_ | _ALIQC[Carbamidomethyl (C)]AK_ | _[Acetyl (Protein N-term)]MEAFQELR_ |
| PEAKS | AM(+15.99)DVYQK | ALIQC(+57.02)AK | M(+42.01)EAFQELR |
| i2MassChroQ | AM[MOD:00719]DVYQK | ALIQC[MOD:00397]AK | not reported |
| FragPipe | AM[15.9949]DVYQK | ALIQC[57.0215]AK | n[42.0106]MEAFQELR |
| WOMBAT | AM[Oxidation]DVYQK | ALIQC[Carbamidomethyl]AK | [Acetyl]-MEAFQELR |
| AlphaPept | AoxMDVYQK | ALIQcCAK | aMEAFQELR |
| MS-Angel | AMDVYQK; Oxidation (M2) | ALIQCAK; Carbamidomethyl (C5) | MEAFQELR; Acetyl (Protein N-term) |
| Proline | AMDVYQK; Oxidation (M2) | ALIQCAK; Carbamidomethyl (C5) | MEAFQELR; Acetyl (Protein N-term) |
| quantms | AMDVYQK only the unmodified form reported | ALIQC(Carbamidomethyl)AK | not reported |
| **APB parsing rules**<br>modification maps | 7 notations → UNIMOD:35 | 7 notations → UNIMOD:4 | 7 notations → UNIMOD:1 |
| **ProForma 2.0** | AM[UNIMOD:35]DVYQK | ALIQC[UNIMOD:4]AK | [UNIMOD:1]-MEAFQELR |

## B

##### As written in the parameter file, and as APB reads it

| | Precursor tolerance<br>raw line (gray) → typed value (blue) | Fragment tolerance | Enzyme | Normalization |
|---|---|---|---|---|
| DIA-NN | --mass-acc-ms1 15<br>→ 15 ppm | --mass-acc 20<br>→ 20 ppm | --cut K*,R*<br>→ Trypsin/P | Normalisation disabled<br>not read yet |
| MaxQuant | <mainSearchTol>4.5</mainSearchTol><br>→ 4.5 ppm | <MatchTolerance>20</MatchTolerance><br><MatchToleranceInPpm>True<br></MatchToleranceInPpm><br>→ 20 ppm | <enzymes> <string>Trypsin/P</string><br>→ Trypsin/P | <lfqNormType>1</lfqNormType><br>not read yet |
| FragPipe | msfragger.precursor_mass_lower=-20<br>msfragger.precursor_mass_upper=20<br>msfragger.precursor_mass_units=1<br>→ 20 ppm | msfragger.fragment_mass_tolerance=<br>0.02 msfragger.fragment_mass_units=0<br>→ 0.02 Da | msfragger.search_enzyme_name_1=<br>stricttrypsin<br>→ Trypsin/P | ionquant.normalization=1<br>not read yet |
| Sage | "precursor_tol": {"ppm": [-20.0,<br>20.0]}<br>→ 20 ppm | "fragment_tol": {"ppm": [-20.0,<br>20.0]}<br>→ 20 ppm | "cleave_at": "KR"<br>→ Trypsin/P | not in the file |
| Spectronaut | Main Search: Dynamic<br>→ automatic | Main Search: Dynamic<br>→ automatic | Enzymes / Cleavage Rules: Trypsin/P<br>→ Trypsin/P | Cross-Run Normalization: True<br>→ True |
| AlphaDIA | target_ms1_tolerance: 10<br>→ 10 ppm | target_ms2_tolerance: 15<br>→ 15 ppm | enzyme: trypsin/p<br>→ Trypsin/P | normalize_lfq: True<br>not read yet |
| PEAKS | Precursor Mass Error Tolerance:<br>10.00ppm<br>→ 10 ppm | Fragment Mass Error Tolerance:<br>0.02Da<br>→ 0.02 Da | Enzyme: Trypsin<br>→ Trypsin | Normalization Method: No<br>Normalization<br>→ No Normalization |
| MetaMorpheus | PrecursorMassTolerance = "±10.0000<br>PPM"<br>→ 10 ppm | ProductMassTolerance = "±20.0000<br>PPM"<br>→ 20 ppm | Protease = "trypsin"<br>→ Trypsin | Normalize = false<br>→ False |
| **APB typed**<br>parameter record | **precursor_mass_tolerance**<br>value with unit (ppm \| Da) or automatic | **fragment_mass_tolerance** | **enzyme**<br>one name per cleavage rule | **abundance_normalization_ions**<br>the software's own term |

Figure S3: Examples of heterogeneity in peptidoform and parameter notations. (A) Three peptidoforms as the tools write them: oxidized methionine, carbamidomethylated cysteine, and protein N-terminal acetylation. (B) Settings as each tool writes them in its parameter file (gray, monospaced) and the value, with its unit where it has one, that APB records (below); settings that APB does not read yet are marked.

ered for 98–100% of submissions, and the software version for 95%. Most gaps lie in the parameter files. The ten submissions without a version are `PEAKS` submissions whose files state none. The `DIA-NN`, `MS-Angel`, and `Proline` files and 12 of 17 `PEAKS` files state no protein FDR. One gap lies in APB: the parser does not read the mass tolerances that all four `Spectronaut` 21 files state.

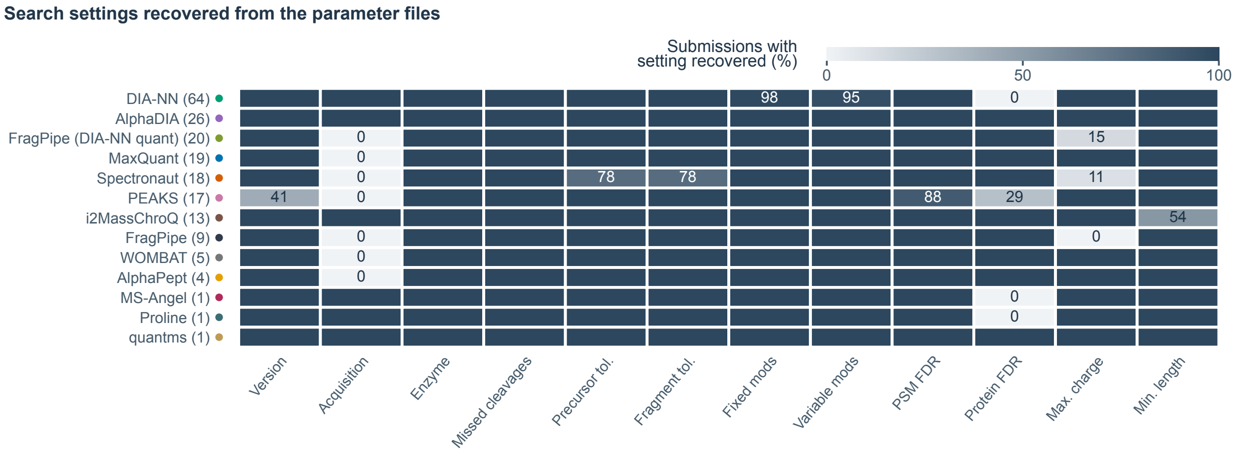


Figure S4: Settings recovered from each submitted parameter file, as the share of submissions per quantification tool, with the number of submissions in parentheses; numbers mark cells below 100%. Acquisition mode counts when the record states DDA or DIA. Automatic mass tolerances count as recovered; modifications count when the record lists at least one.

The acquisition mode is known for 54% of submissions. `DIA-NN` logs mark a DDA search with `--dda`, and the parser records DIA otherwise. The `quantms` parser reads the mode from the pipeline settings. The `AlphaDIA` parser always records DIA, and the `i2MassChroQ`, `MS-Angel`, and `Proline` parsers always record DDA. For the other quantification tools, the mode stays unknown.

### S9 Catalog

`apb-catalog` keeps its terms in sets, one per topic, such as `abundance_alpha`, `confidence_alpha`, and `chromatography_alpha`; the suffix `_alpha` marks a set as a draft, as `_beta` would mark a later one. A further set, `apb_alpha`, names the columns that apb2 writes for every quantification tool, such as ProForma_ion, and the columns that carry a role; obs_annotation_alpha, fasta_peptides_alpha, and fasta_proteins_alpha name the columns that apb2 annotate and apb-fasta add. All sets share the structure of Table S6; Figure S5 shows how its tables relate. Each set names its terms by key: a key stands for a kind and the qualifier values that it fixes, so intensity_norm_settings is an intensity with the cross-run normalization set in the search. Each set also has a prefix that limits a lookup to it: a for abundance_alpha, c for confidence_alpha, rt for chromatography_alpha, apb for apb_alpha, annot for obs_annotation_alpha, and fpep and fprot for the two FASTA sets; the prefix v asks the quantification tool's own names alone. The resolver DataCatalog reads a result through the sets that it names; apb_alpha answers only when named. It answers a key with the layer or feature-table (var) column that the key maps to for the tool and version of the result, or with none when the tool records none. It raises UnresolvedField when the answer is ambiguous or unknown. Catalog answers the same request from a tool name and version, without a result. The apb-catalog documentation describes the interface.

#### S9.1 Abundance set

The abundance_alpha set gives each abundance layer a kind, intensity or ibaq, and three qualifiers: normalization, aggregation, and MS level. The normalization is none, run_setting, or unknown; run_setting means the cross-run normalization set in the search, which may be none. The aggregation states how a value combines lower-level features: none for a feature's own signal, sum, top_n, maxlfq, or directlfq. The MS level is ms1 or ms2. The keys intensity and intensity_ms1 fix the normalization none, intensity_norm_settings and intensity_ms1_norm_settings fix run_setting, and intensity_norm_unknown fixes unknown; the two ms1 keys also fix the MS level. The key ibaq fixes the kind alone. apb-catalog meaning abundance_alpha prints the whole set, and the

Table S6: Structure shared by all sets of the catalog, which map each term to the field of every quantification tool that holds it. Purple rows describe a term, as the application developer states it in `terms.csv`; an application asks for the term by its key. The other rows describe the entry that the integrator writes in `catalog.csv` for each quantification tool and version, with `DIA-NN` and `AlphaDIA` as examples; the blue row is the name of the field returned. Bold marks the words the main text uses for these columns. The requirement field indicates how essential the term is to the application.

| Column | Holds | Examples |
|---|---|---|
| **set** | The set of the **catalog** that the term belongs to, one per topic | abundance_alpha, confidence_alpha, chromatography_alpha, apb_alpha |
| **key** | The **term**'s name, which an application asks for; it fixes the kind and the qualifiers | intensity, intensity_norm_settings, q_value, retention_time |
| **location** | The level and slot that hold the term: a layer holds one value per run, a var column one per feature | peptidoform_ion/layers, protein/var, */obs (every level) |
| **kind** | What the value measures; each set defines its own kinds | intensity, ibaq, q_value, pep, retention_time |
| **qualifier, value** | Further properties, each as qualifier = value, which the key fixes | intensity: normalization = none; intensity_norm_settings: normalization = run_setting |
| **requirement** | How strongly applications need the term: required, recommended, or optional | With enforce = required, a missing required field stops the catalog from opening; a missing recommended field, such as unnormalized intensity in AlphaDIA, logs a warning. With enforce = recommended, both stop |
| **definition** | What the term means, in words | unnormalized ion intensity |
| **cv_accession** | Controlled-vocabulary accessions for the term's meaning, such as PSI-MS terms; may be empty | intensity: MS:1001840 |
| **source** | The quantification tool and the version label of the entry, as tool/version; a label matches a pattern of version strings | DIA-NN/1.8–1.9, DIA-NN/* (every version), AlphaDIA/2.x |
| **name** | The **field**'s name: the layer or column that answers the term, as layer/column | DIA-NN: Precursor_Quantity/-, -/Global_Q_Value; AlphaDIA 2.x: precursor_intensity/- |
| **fixed** | Whether the key fixes the value of this qualifier, as the term states it | intensity_norm_settings, normalization: true |
| **basis, evidence_url** | Why the field answers the term, as free text, with an optional source URL | DIA-NN, Precursor_Quantity: "Quantity means non-normalised quantity" (DIA-NN output reference); source: github.com/vdemichev/DiaNN |

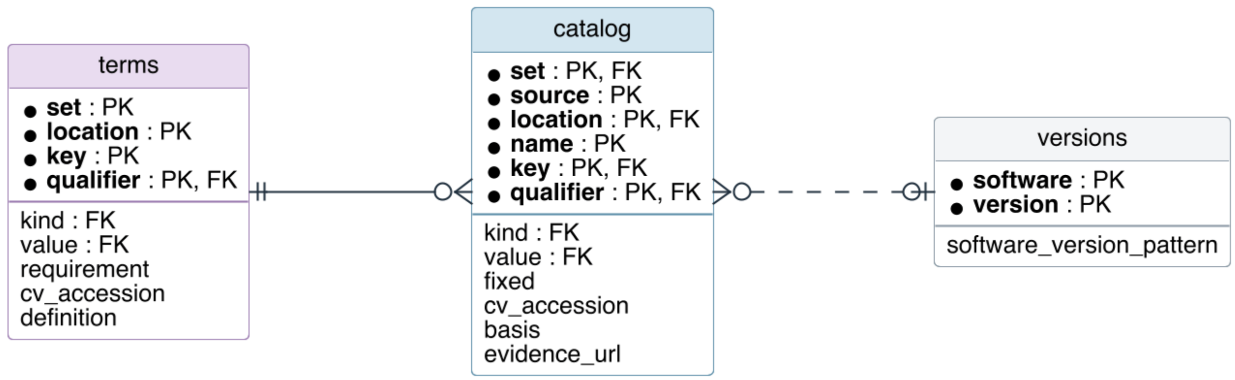


Figure S5: Entity-relationship diagram of the catalog tables `terms`, `catalog`, and `versions`, with the primary keys (PK, the columns above the line) and foreign keys (FK) that the catalog's table schema declares. Line ends mark exactly one (double bar), zero or one (ring and bar), and zero or more (ring and crow's foot). Each term key, the set, location, and key of `terms`, has zero or more catalog entries, one for each tool version and qualifier; the solid line is the declared foreign key. The dashed line links an entry's `source`, such as `DIA-NN/2.x`, to at most one version label; `DIA-NN/*` matches every version. The sets, kinds, qualifiers, and qualifier values, which the remaining foreign keys reference, are not shown.

catalog viewer (`https://fgcz-ms.uzh.ch/public/apb-catalog/`) shows, for each quantification tool, version, and level, the layer that holds each abundance.

#### S9.2 Confidence set

The `confidence_alpha` set gives each posterior error probability or q-value a kind, `pep`, `q_value`, or `library_q_value`, and a stage, identification or quantification. The keys `pep`, `q_value`, and `library_q_value` fix the identification stage; `quantification_q_value` is a q-value of the quantification, such as `Sage`'s q-value of the MS1 peak. Per-run values are layers. A tool reports some confidences once per feature for all runs. These are feature-table columns, which `var` returns, such as `Spectronaut`'s experiment-wide protein q-value `PG_Qvalue`. The key `q_value` thus names the per-run q-value in the layers and the experiment-wide q-value in the feature table, and the slot a consumer reads decides which it gets: `layers["q_value"]` returns the per-run layer, `var["q_value"]` the feature-table column.

#### S9.3 Reading a result through the catalog

DataCatalog gives each level the interface of an AnnData object[29]: the primary abundance X, the further quantity layers layers, the feature table var, the sample table obs, and the names obs_names and var_names. As in AnnData, X and the layers are NumPy matrices of samples by features[30]; the columns of var and obs are Polars series[31]. A name in layers, var, or obs is looked up as a key first and as the tool's own name second, as cf_xarray looks up the variables of a dataset by their Climate and Forecast (CF) standard names first[32]. The tool's columns thus stay readable under names taken from their headers, even where no set covers them. A tool may name a column as a key is named, as Sage names one q_value; a prefix then makes the lookup exact: c:q_value asks the confidence set alone, and v:q_value the tool's own column alone. For the DIA-NN 1.9.2 report of the export example (Section S11.2):

```
from pathlib import Path

from apb2.api import read_parsed_levels
from apb_catalog.api import Catalog, DataCatalog

# result.h5mu: an APB result written by apb2 convert
result = read_parsed_levels(Path("result.h5mu"))
catalog = DataCatalog(result, "abundance_alpha", "confidence_alpha", "apb_alpha")
ion = catalog.levels["peptidoform_ion"]  # AnnData's slots: X, layers, var, obs

ion.layers                             # key -> NumPy matrix, runs by precursors
ion.layers.keys()                      # intensity, intensity_norm_settings, intensity_ms1,
                                       # intensity_ms1_norm_settings, q_value, pep
ion.layers["intensity"]                # Precursor_Quantity: 6 runs by 97,974 precursors
ion.layers["intensity_norm_settings"]  # Precursor_Normalised, the primary abundance X
ion.layers["q_value"]                  # Q_Value: one per run and precursor
ion.var                                # key -> Polars Series, one value per precursor
ion.var["q_value"]                     # Global_Q_Value: one per precursor
ion.var["c:q_value"]                   # the same, from the confidence set alone
ion.var["peptidoform_ion"]             # ProForma_ion, from the apb_alpha set
ion.layers["Quantity_Quality"]         # no key: DIA-NN's own layer of that name
ion.layers["v:Q_Value"]                # DIA-NN's own layer alone, never a key
ion.layers["intensities"]              # KeyError: neither a key nor a layer name

diann = Catalog("DIA-NN", "1.9.2", "abundance_alpha")      # no result needed
diann.levels["peptidoform_ion"].layers["intensity"]  # 'Precursor_Quantity', a name
```

`Catalog` answers the same keys from the name and version of a quantification tool alone: its levels have the same slots and hold the names of the layers and columns, which `apb-export` records in the JSON summary of an export (Section S11.3). The same lines read the result of any quantification tool. At the protein level of the example, two layers carry the normalization set in the search, `PG_Normalised` and `PG_MaxLFQ`, so `layers["intensity_norm_settings"]` raises `UnresolvedField`, and `layers.get("intensity_norm_settings", aggregation="maxlfq")` returns the `PG_MaxLFQ` layer. For an `AlphaDIA` result, which holds no intensity without normalization, `layers["intensity"]` raises `KeyError`, as AnnData does for a missing layer, and `layers.get("intensity")` returns `None`.

### S10 ProteoBench scoring example

A `ProteoBench` submission is the result table and parameter file of one run of a quantification tool on a module's raw data. `ProteoBench` scores it by how well the observed log2 fold changes match the expected species ratios[33–35]. `apb-proteobench` computes these scores from a ProteoBridge result. It can also write `ProteoBench`'s intermediate output, from which `pmultiqc` builds a quality-control report[36].

#### S10.1 Data and commands

The example is submission `269bb831` to the DIA Astral module (`dia_astral`), processed with `DIA-NN` 2.5.0 Academia. Its archive holds the report (`input_file.parquet`, 59 MB), the `DIA-NN` log and `ProteoBench`'s own per-feature result. The module FASTA holds the human, yeast, and *E. coli* proteomes and contaminants, 31,889 proteins in total.

```
H=269bb8310dc3f501834aaab5ca6fe72791c426e3
U=https://proteobench.cubimed.rub.de
curl -O $U/datasets/$H/${H}_data.zip
curl -O $U/fasta/ProteoBenchFASTA_MixedSpecies_HYE.zip
unzip ${H}_data.zip
unzip ProteoBenchFASTA_MixedSpecies_HYE.zip
mv input_file.parquet report.parquet
mv param_0..txt report.log.txt
uv venv --python 3.13 && source .venv/bin/activate
uv pip install apb2==0.1.4 apb-catalog==0.1.6 apb-fasta==0.1.2 \
  apb-proteobench==0.2.0 protein-fasta==0.3.2 prozor==0.1.2
```

Three commands convert the report, match its peptides against the module FASTA, and score the result. Each reads the result of the previous step, so no step rereads the software files. The option `--format` writes Parquet or DuckDB instead of HDF5.

```
apb2 convert report.parquet --params report.log.txt \
  --format hdf5 --output converted
apb-fasta verify-peptides converted.h5mu \
  ProteoBenchFASTA_MixedSpecies_HYE.fasta \
  --output fasta-checked.h5mu
apb-proteobench benchmark fasta-checked.h5mu \
  dia_astral scored.h5mu
```

#### S10.2 Outputs of each step

- `apb2 convert`: the peptidoform-ion level holds 6 samples × 113,446 ions in 34 layers, and the protein level 6 samples × 11,083 protein groups in 5 layers. Values that `DIA-NN` repeats in every run, such as the library and global q-values, become feature-table columns. Without gene annotations, `DIA-NN` reports zero for every gene quantity, so the result leaves out `Genes_MaxLFQ` as effectively empty.
- `apb-fasta verify-peptides` finds the peptides of all 113,446 ions in the FASTA, with no decoys, and records the FASTA checksum.
- `apb-proteobench benchmark` records the module definition, the expected ratios, and the `ProteoBench` code revision it reproduces. It scores the primary abundance `Precursor_Normalised` for completeness cutoffs of one to six runs and stores per-feature diagnostics in `varm`. At cutoff 1, it scores 111,489 ions, with a global median absolute error of 0.218.

#### S10.3 Scoring criteria

`apb-fasta` matches modification-stripped peptide sequences through `prozor`[37,38], using the Aho-Corasick algorithm[39]. `apb-proteobench` scores an ion that is observed, is neither a contaminant nor a decoy, and matches exactly one species. At completeness cutoff $n$, it scores the ions observed in at least $n$ runs. The command scores the primary abundance. For a module that requires a normalization, it scores the layer that the abundance set names.

### S11 Exports

#### S11.1 Export rules

An export rule is a JSON file that names its export target and the target versions it supports. Each table has a base for all levels and blocks for single levels, which override the base. Each layer and column pairs a `name`, the field of the application, with a `source`: a role, a column that `apb2` builds, such as `ProForma_ion`, or a catalog term. A new export target, or a new version of a target, needs a new export rule. The apb-export documentation describes each export target.

In Excerpt S1, the intensity comes from the abundance set, with the primary abundance as fallback. `protein_Id` and `precursor_Id` come from the `apb_alpha` set: the column with the `protein_assignment` role and the `ProForma_ion` identifier. The export uses the first listed level that exists in the result and fits its quantification tool. The `software` entry limits the peptide level to `MaxQuant` results.

**Excerpt S1.** `prolfqua` export rule, format version 2.11: an intensity layer with a fallback, level selection by quantification tool, and two feature columns. From `prolfqua/v2_11/rules.json` in `apb-export` commit `8885160`.

```
{
  "target_name": "prolfquapp",
  "target_version_pattern": "^2\\.11\\.",
  "tables": [
    {
      "base": {
        "measurements": {
          "layers": [
            {
              "name": "intensity",
              "source": {"catalog": "abundance_alpha", "layers": "intensity"},
              "fallback": [{"measurements": "primary_layer"}]
            }
          ]
        },
        "columns": {
          "var": [
```

```
            {
              "name": "protein_Id",
              "source": {"catalog": "apb_alpha", "var": "protein_assignment"},
              "drop_missing": true
            }
          ]
        }
      },
      "levels": {
        "peptide": {"software": ["MaxQuant"]},
        "peptidoform_ion": {
          "columns": {
            "var": [
              {
                "name": "precursor_Id",
                "source": {"catalog": "apb_alpha", "var": "peptidoform_ion"}
              }
            ]
          }
        }
      }
    }
  ]
}
```

#### S11.2 Reproducing the export example

The export example uses `ProteoBench` submission `0132a468` to the DIA AIF module, a `DIA-NN` 1.9.2 report of six runs. The commands below download it, the module FASTA, and the module SDRF, which ships with `apb-proteobench`, and install `apb-export` from PyPI with `uv`. The versions are the releases with which we verified the example on 23

November 2026.

```
H=0132a468ec6ae51a527d82560ff4a40f832f975c
U=https://proteobench.cubimed.rub.de
curl -O $U/datasets/$H/${H}_data.zip
curl -O $U/fasta/ProteoBenchFASTA_MixedSpecies_HYE.zip
unzip ${H}_data.zip
unzip ProteoBenchFASTA_MixedSpecies_HYE.zip
mv input_file.txt report.tsv
mv param_0..txt report.log.txt
R=https://raw.githubusercontent.com/anndata-omics-bridge/apb-proteobench/v0.2.0
curl -O $R/src/apb_proteobench/data/modules/dia_aif.sdrf.tsv
uv venv --python 3.13 && source .venv/bin/activate
uv pip install apb2==0.1.4 apb-catalog==0.1.6 apb-fasta==0.1.2 \
  apb-export==0.2.3 protein-fasta==0.3.2 prozor==0.1.2
```

The `apb-export` command of the main text then writes `prolfqua.h5ad`. From Python, the same steps run in memory, and `Exporter` writes the same file. `exporter.levels` limits the conversion to the levels that the export rule reads.

```
from pathlib import Path

from apb2.api import AnnotationCompiler, ParseRuleCompiler
from apb_export.api import Exporter
from apb_fasta.api import FastaAnnotator

exporter = Exporter("prolfqua")
parsed = ParseRuleCompiler.detect(
    Path("report.tsv"), Path("report.log.txt"), requested_levels=exporter.levels
).compile().parse()
parsed = AnnotationCompiler().compile(Path("dia_aif.sdrf.tsv")).parse(parsed).annotate().parsed
fasta = FastaAnnotator.read([Path("ProteoBenchFASTA_MixedSpecies_HYE.fasta")])
parsed = fasta.verify_peptides(parsed).parsed
parsed = fasta.merge_annotations(parsed).parsed
exporter.write(exporter.export(parsed), Path("prolfqua.h5ad"))
```

#### S11.3 Catalog choices of the prolfqua export

The `prolfqua` export rule requests an intensity without normalization. Where the abundance set names none, the export takes the primary abundance (Excerpt S1), logs a warning and records the choice in its JSON summary. Confidence values come from the confidence set; experiment-wide values come from feature-table columns and repeat in every run. For the example, the export chose intensity=Precursor_Quantity, qValue=Q_Value, pg_qValue=PG_Q_Value, pg_qValue_experiment=Lib_PG_Q_Value, and pep=PEP.

#### S11.4 Export checks

The apb-export repository currently checks nine of its ten export rules; the exploreDE rule has no check yet. Each check exports a synthetic DIA-NN report and opens the file with the application's own reader. msmu reads the wide psm modality, still experimental, with read_h5mu; its filtering, normalization, and aggregation give the same peptide and protein values as from its own DIA-NN reader[40]. prolfquapp's LFQData_from_anndata rebuilds the LFQData object, with its protein, peptide, and precursor hierarchy and its sample factors[41,42]. Its preprocess_APB reader gives the same peptide values, and medpolish aggregation the same protein values, as its own DIA-NN reader preprocess_DIANN. ProteoPy reads peptides as its primary input. The proteopy_ion export writes one feature per precursor and gives the same precursors, values, and protein identifiers as ProteoPy's own DIA-NN reader at precursor level; check_proteodata accepts this file as peptide data and the proteopy export as protein data[43]. AlphaPeptTools' mulink_from_anndatas links the exported modalities into one MuData object[44]. MSstats' dataProcess summarizes the long table to proteins, and groupComparison compares the two conditions[45]. QFeatures reads the MuData file with readH5MU, links the levels, and aggregates precursors to proteins[46]. perseuspy reads the Perseus table as Perseus reads its own files, with typed sample columns. The QPX export is validated against the QPX schemas and read by pmultiqc[36,47]. Continuous integration runs the msmu, ProteoPy, AlphaPeptTools, and Perseus checks in one container image, and the QFeatures and MSstats checks in an R image.

#### S12 Export web app

The ProteoBridge export web app (main-text Figure 5B) converts one result without installation. The user chooses the quantification tool and uploads its result files. Optional uploads are the parameter file, a sample table in SDRF-Proteomics or `prolfquapp` format, and a FASTA database; only `Sage` requires the parameter file. Uploads may total 2.1 GB, and example inputs run without an upload. The user then chooses an export target or the result itself as AnnData/MuData, Parquet, or DuckDB. The page offers the downloads with quality-control summaries of the exported matrices. The app is `apb-export-ui`. The command `uv tool install --python 3.13 apb-export-ui==0.3.0` installs it from PyPI, and `apb-export-web start` opens it in the browser. Launchers in its repository do both on macOS, Linux, and Windows and install `uv` if it is missing.

(13) Demichev, V.; Messner, C. B.; Vernardis, S. I.; Lilley, K. S.; Ralser, M. DIA-NN: neural

networks and interference correction enable deep proteome coverage in high throughput. *Nature Methods* **2020**, *17*, 41–44, DOI: 10.1038/s41592-019-0638-x.
